# Cancer-specific association between Tau (*MAPT*) and cellular pathways, clinical outcome, and drug response

**DOI:** 10.1101/2023.07.04.547660

**Authors:** Maurizio Callari, Martina Sola, Claudia Magrin, Andrea Rinaldi, Marco Bolis, Paolo Paganetti, Luca Colnaghi, Stéphanie Papin

## Abstract

Tau (*MAPT*) is a microtubule-associated protein causing common neurodegenerative diseases or inherited frontotemporal lobar degenerations. Emerging evidence for non-canonical functions of Tau in DNA repair and P53 regulation suggests its involvement in cancer. Indeed, preliminary studies have correlated Tau expression with cancer survival or response to therapies. To bring new evidence for a relevant role of Tau in cancer, we carried out an *in silico* pan-cancer analysis of *MAPT* transcriptomic profile in over 10000 clinical samples from 32 cancer types and over 1300 pre-clinical samples from 28 cancer types provided by the TCGA and the DEPMAP datasets respectively. *MAPT* expression associated with key cancer hallmarks including inflammation, proliferation, and epithelial to mesenchymal transition, showing cancer-specific patterns. In some cancer types, *MAPT* functional networks were affected by P53 mutational status. We identified new associations of *MAPT* with clinical outcomes and drug response in a context-specific manner. Overall, our findings indicate that the *MAPT* gene is a potential major player in multiple types of cancer. Importantly, the impact of Tau on cancer seems to be heavily influenced by the specific cellular environment.

## Introduction

It has been known for decades that the microtubule-binding protein Tau plays a role in causing debilitating neurodegenerative disorders(1). Indeed, various rare autosomal dominant mutations in the *MAPT* gene encoding for the protein Tau cause frontotemporal lobar degeneration with Tau pathology (FTLD-Tau), which defines a small group of progressive frontotemporal lobar degenerations(2, 3). FTLD-Tau, together with Alzheimer’s disease (AD), belongs to a group of neurodegenerative disorders that are identified by the presence of an abnormal fibrillar form of hyperphosphorylated Tau protein inside the cells(4, 5). Subcellular distribution and activities of Tau that may not be related to its association with microtubules were recently described, including functions linking Tau to nucleic acids(6). For instance, studies have reported the accumulation of DNA lesions in the brains of *MAPT* knock-out (*MAPT* KO) mice and primary neurons(7, 8). Additionally, loss of heterochromatin has been observed in the brains of individuals with AD and in animal models of tauopathies(9–11). Supporting this, Tau is present in the cell nucleus with a specific phosphorylation pattern(12) and binds to DNA in a sequence-independent manner(11, 13, 14). Also, Tau depletion increases the sensitivity to DNA-damaging drugs in a xenograft model of breast cancer(15). Moreover, Tau depletion in cells recovering from acute DNA damage resulted in reduced initiation of programmed cell death because of P53 destabilization; an effect that was compensated by increased cellular senescence induction(16).

These findings suggest a molecular connection between P53 and Tau and a potential involvement in cancer(17), creating a possible link between neurodegeneration and cancer, two prevalent age-related human diseases(18). Tau expression has been correlated with the response to microtubule targeting drugs and other cancer treatments(17). In the central nervous system (CNS) cancers neuroblastoma and glioma, Tau level correlated with survival(19, 20). Increased Tau protein was also associated with Isocitrate Dehydrogenase (*IDH1*) mutations in glioma (a likely driver of formation and development of this type of cancer(21)) and with improved prognosis and response to therapy. However, the comprehensive role of Tau in neoplastic conditions is still unclear.

Considering these initial reports supporting possible roles of Tau in cancer, we report herein the outcome of a set of pan-cancer analyses aimed at demonstrating the relevance of Tau in malignancies, identifying the genes and pathways associated with Tau, quantifying the impact of Tau expression on the clinical outcome and drug response, and exploring the possible interplay with P53. We mined the pan-cancer TCGA(22) dataset as well as the DEPMAP(23) resource, for a total of over 10000 clinical samples and over 1300 pre- clinical samples. Altogether, the obtained results support a critical and context-dependent role of Tau in cancer.

## Results

### *MAPT* co-expression analysis highlighted cancer-specific associations

To shed light on the relevance of Tau in cancer, we evaluated *MAPT* gene expression values in 32 distinct cancer types mining the TCGA pan-cancer cohort. *MAPT* expression was highly variable, with brain glioblastoma multiforme (GBM), lower grade glioma (LGG), and neuroendocrine (pheochromocytoma and paraganglioma (PCPG) tumors showing the highest expression followed by breast cancer (BRCA) (**Figure 1a**).

**Figure 1.**
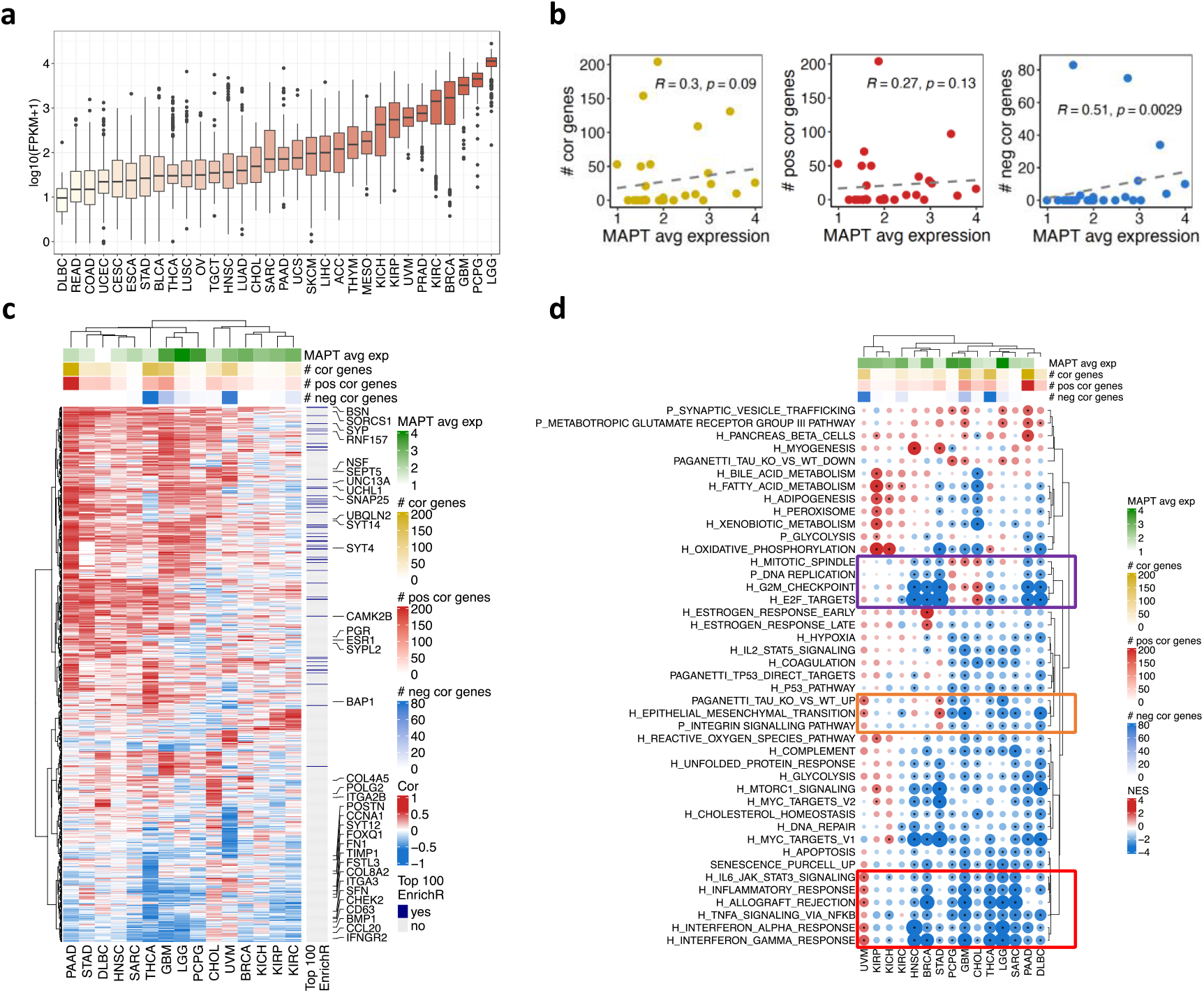
- Pan-cancer evaluation of MAPT expression and transcriptional associations. **(a)** MAPT expression in the TCGA according to cancer type; **(b)** association between MAPT average expression and the number of correlated genes per cancer type. The total number (left), the number of positively correlated genes (centre), and the number of negatively correlated genes (right) were evaluated. **(c)** Heatmap of all genes correlated with MAPT expression in at least one cancer type. Both genes and cancer types are ordered based on the hierarchical clustering of the correlation values. Genes in the list of top 100 genes co-expressed with MAPT according to EnrichR (ARCHS^4^ dataset) are indicated. Selected genes are highlighted. For each cancer type, the average MAPT expression and the number of more than one gene correlated with MAPT are shown. The whole pan-cancer results are reported in Figure S1. **(d)** GeneSet Enrichment Analysis on the genes ranked accordingly to their correlation with MAPT expression in each cancer type. A negative Normalised Enrichment Score (NES) means down-regulation of the geneset for high MAPT expression and vice versa for positive NES. Colored boxes indicate genesets commented in the text.

*MAPT* expression was next correlated with all expressed genes in each cancer type. The number of genes displaying a positive or negative correlation with *MAPT* expression was highly variable across cancers. We found correlated genes (absolute correlation threshold = 0.6) in 15 out of 32 cancer types. The highest numbers of correlated genes were observed in pancreatic adenocarcinoma (PAAD), thyroid carcinoma (THCA), glioblastoma (GBM), and uveal melanoma (UVM) (**Figure S1a**). In some cases (e.g. PAAD), positive correlations were predominant, while negative correlations prevailed in THCA and UVM. We checked whether the total number of genes correlating with *MAPT* or the number of genes with a positive or a negative correlation depended on *MAPT* expression levels. Only a low-moderate correlation was observed in all cases (**Figure 1b**), suggesting only a weak link between the expression levels and the potential relevance of *MAPT* within cancer biological networks.

**Figure S1.**
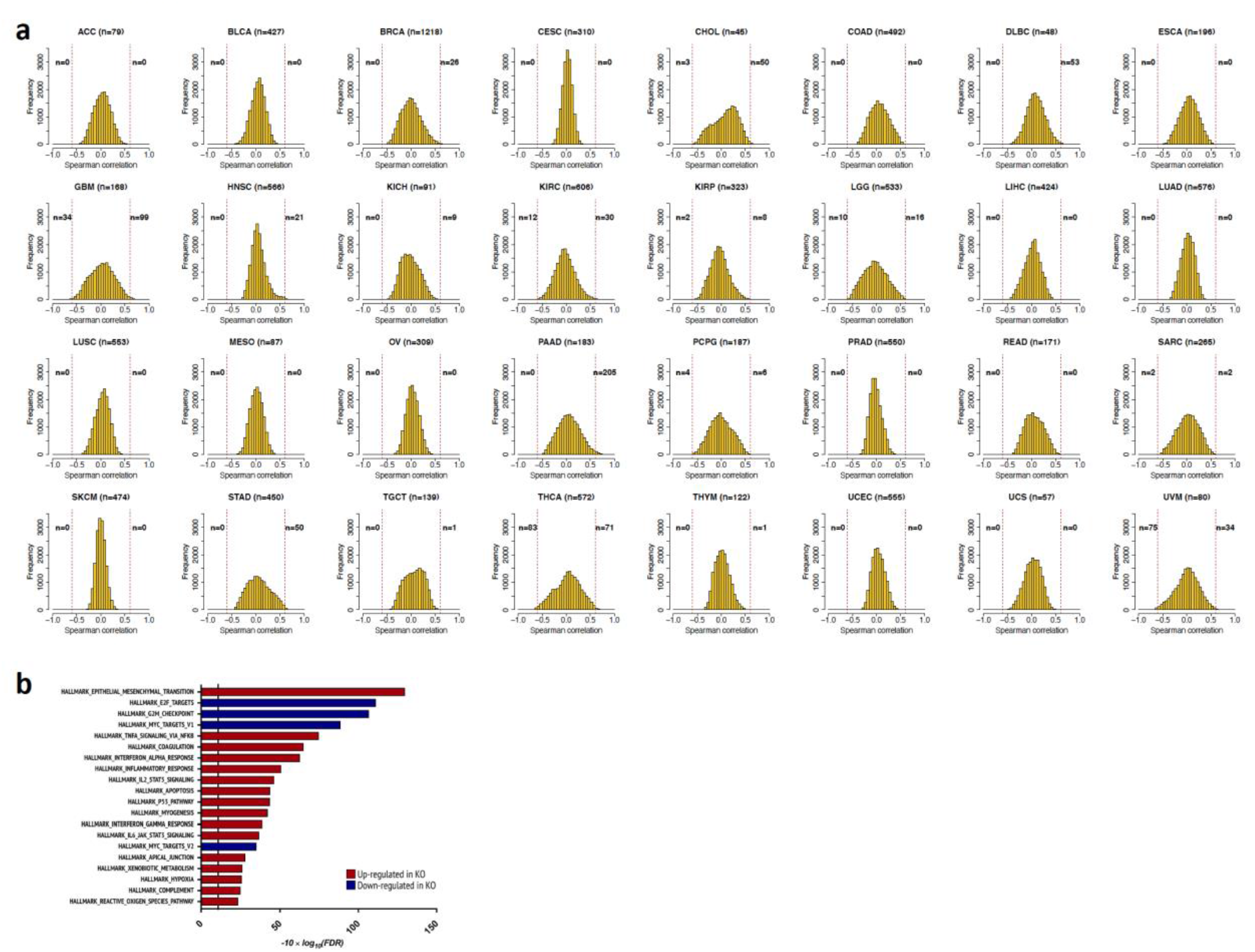
**- (a)** Correlation of MAPT gene expression with all expressed genes. Each histogram shows the correlation value distribution obtained for each cancer type. The number of genes with high positive/negative correlation (i.e. above 0.6 or below -0.6) is reported. **(b)** Top 20 enriched Hallmark genesets in the comparison between MAPT KO and WT neuroblastoma cells.

From the *MAPT*-all gene correlation analysis, we identified 809 genes with a significant correlation with *MAPT* in at least one cancer type (**Figure 1c** and **Figure S2**). The unsupervised analysis of their correlation patterns grouped the tumor types according to their organ or apparatus of origin. Indeed, ESCA, STAT, COAD, and READ clustered together, as well as GBM, LGG, and PCPG or KICH, KIRP, and KIRC (**Figure S2**). We intersected our list of genes with the top 100 genes co-expressed with *MAPT* according to EnrichR, which uses the ARCHS^4^ dataset (see Methods). Fifty-two genes were included in the 809 identified in the TCGA (**Figure 1c** and **Figure S2**). Considering that any kind of human tissue is included in ARCHS^4^ and that the pan-cancer correlation highlighted substantial differences among cancer types, the overlap was remarkable.

In the 15 cancer types with significantly correlated genes, the correlation analysis was complemented with a GeneSet Enrichment Analysis (GSEA). For each cancer type, genes ranked by correlation with *MAPT* became the input of a GSEA to identify biological processes and pathways having a positive or negative correlation with *MAPT* expression (**Figure 1d**). We evaluated the Cancer Hallmark collection (24), the PANTHER Pathway collection (25), and additional custom genesets. Indeed, in light of our previous findings showing that Tau stabilizes P53 and negatively modulates senescence in response to DNA damage(16), we include four additional genesets capturing senescence(26–29) or direct P53 target genes (30–33). We also included a geneset containing the top 100 genes dysregulated in *MAPT* KO neuroblastoma cells (developed by us and previously described (16)) when compared to Tau-expressing neuroblastoma cells (**Figure S1b**).

Biological interpretation of the results highlighted both commonalities and cancer-specificity in the way *MAPT* expression relates to the tumor ecosystem transcriptomic profile. Genes involved in migration or epithelial to mesenchymal transition (EMT) (*ITGA2B*, *COL4A5*, *FSTL3*, *COL8A2*, *FN1*, *BMP1*, *TIMP1*, *FOXQ1*, *ITGA3*, *POSTN*, *CD63*) correlated with *MAPT* expression positively or negatively depending on the cancer type (**Figure 1c**). Similarly, genesets related to these biological functions (INTEGRIN_SIGNALLING_PATHWAY and EPITHELIAL_MESENCHIMAL_TRANSITION) are also bi-directionally associated with *MAPT* according to the cancer type (**Figure 1d, orange box** and **Figure S3a**). Our signature of upregulated genes in *MAPT* knock-out neuroblastoma cells (PAGANETTI_TAU_KO_VS_WT_UP) (**Figure S1b**) showed an analogous pattern of association. Coherently with the original experiment, a negative enrichment in brain tumors was observed.

An additional pathway showing a strong variation of the NES (normalized enrichment score) according to the cancer type was the OXIDATIVE-PHOSPHORYLATION pathway, which reflects mitochondrial activity. This pathway was negatively enriched in most cancer types with few exceptions: in tumors derived from the kidneys (KICH, KIRP), and to a lesser extent in UVM and THCA, it is strongly positively enriched.

Genesets related to inflammation (INTERFERONα_RESPONSE, INTERFERONγ_RESPONSE, IL6_JAK_ STAT3_SIGNALING, TNF_SIGNALING_VIA_NFKB, INFLAMMATORY_RESPONSE, ALLOGRAFT_REJECTION) globally showed a negative enrichment in all but UVM cancer. The strongest enrichments were found for sarcoma (SARC), GBM, LGG, THCA, and BRCA (**Figure 1d, red box**). IFNGR2 and CCL20 were among the significantly correlated genes related to these biological functions (**Figure 1c**). Negative enrichment was evident for one of the senescence genesets (SENESCENCE_PURCELL_UP (29)), possibly explained by the increase in the secretion of inflammatory mediators during cellular senescence(36), some of which are part of the geneset.

Also, cell cycle-related genesets (G2M_checkpoint, E2F_targets, DNA_REPLICATION, MITOTIC_SPINDLE) were mostly negatively correlated with *MAPT*, in particular in HNSC, BRCA, STAD, PAAD, and DLBC. A few exceptions, with opposite trends, were found for CHOL, PCPG, and GBM (**Figure 1d, violet box, and Figure S3b**). Multiple genes related to DNA replication, repair, and cell cycle progression did show significant correlations, e.g. *CHEK2, SFN*, *POLG2*, *CCNA1*.

Among the 809 genes, we found multiple neuronal genes positively correlating with *MAPT* across most cancer types (e.g. *BSN*, *SORCS1*, *CAMK2B*, *SNAP25*, *SEPT5*, *UNC13A*, *UCHL1*, *NSF*, synaptophysins *SYP* and *SYPL2*, and synaptotagmins *SYT4*, *SYT12* and *SYT14*, some of which have been linked to AD. For example, *SORCS1* polymorphism is associated with AD(34), whereas CAMK2 dysregulation in the hippocampus of AD subjects may contribute to neurofibrillary tangle formation, synaptic degeneration, and memory deficits(35). Consistent with the high number of neuronal genes correlated with *MAPT*, enrichment for the SYNAPTIC_VESICLE_TRAFFICKING geneset was detected, particularly in brain tumors and PAAD, confirming that *MAPT* expression is often associated with a wider activation of a neuronal transcriptional program in cancer. However, this is not ubiquitous across distinct cancer types and a tendency to negative correlation for the same set of genes was observed, for example, in KIRP (**Figure 1c-d** and **Figure S3c**).

Some correlations and enrichment were highly cancer-specific. For example, this is the case for estrogen response genes (e.g. ESTROGEN_RESPONSE_EARLY) specifically enriched in BRCA, with *ESR1* and *PGR* highly correlated with *MAPT* in BRCA. This was consistent with literature data demonstrating that estrogens regulate *MAPT* expression(37–39), supporting the robustness of the present analysis.

Finally, it is worth mentioning that genes involved in ubiquitination processes and linked to cancer were among the significantly correlated genes (*BAP1*, *RNF157*, *UBQLN2*).

In summary, *MAPT* expression levels are associated with major cancer hallmarks, with some commonalities across groups of cancer types but a high degree of tumor specificity, indicating that the biological networks that include *MAPT* are highly context specific.

**Figure S2.**
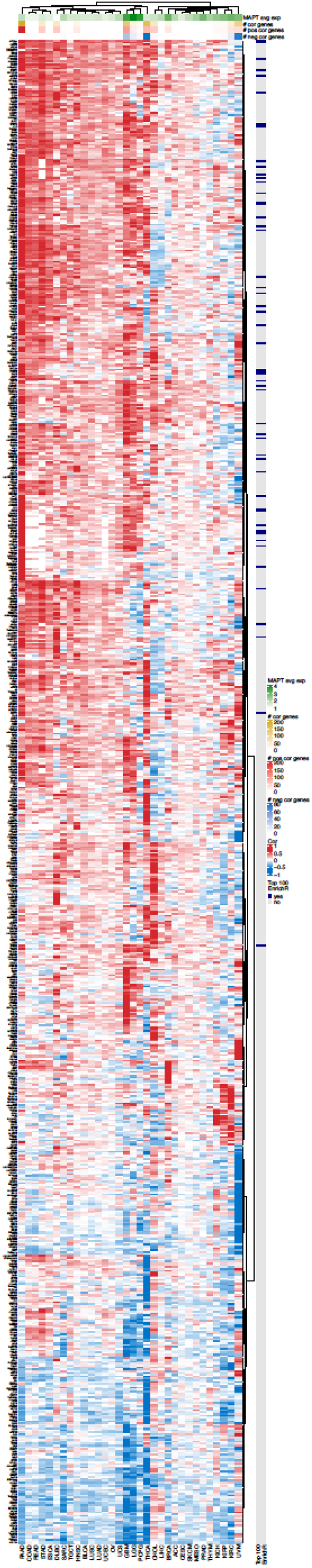
**-** Heatmap of 809 genes correlated with MAPT expression in at least one cancer type. Both genes and cancer types are ordered based on the hierarchical clustering of the correlation values. Fifty-two of the top 100 genes co-expressed with MAPT according to EnrichR (ARCHS^4^ dataset) are overlapping with the 809 genes and are annotated in the heatmap. For each cancer type, the average MAPT expression and the number of genes correlated with MAPT are shown.

**Figure S3.**
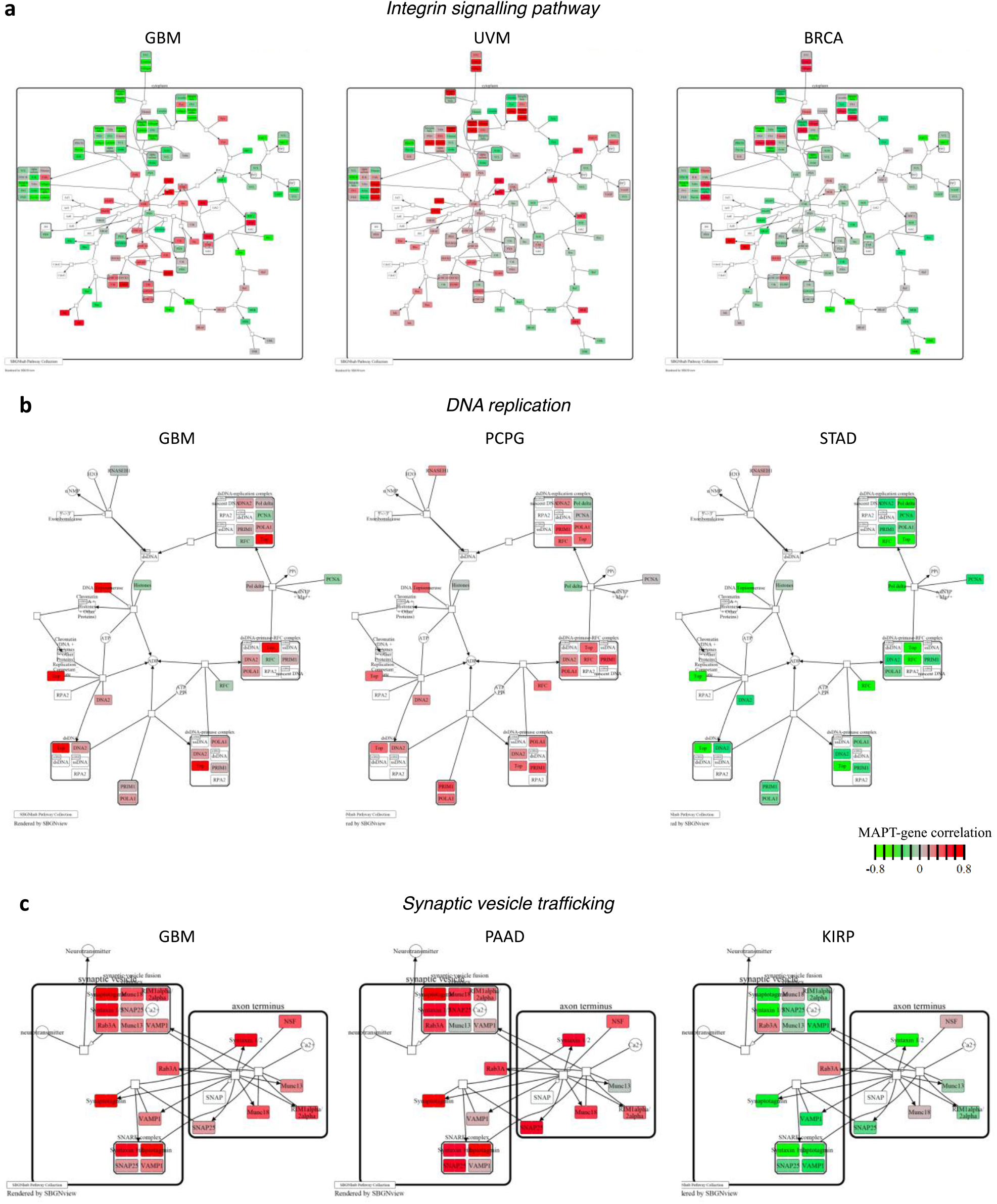
**–** Selected PANTHER pathways in selected cancer types overlaid with the MAPT-gene correlation values for all evaluated genes.

### *MAPT* co-expression analysis highlighted P53-dependent associations

Tau is mostly characterized as a microtubule-stabilizing protein and may influence cancer outcomes through this cellular function. However, we recently described a new role of Tau as a positive modulator of P53 stability in neuroblastoma cells(16). Considering that P53 is mutated in half the tumors, these findings propose a possible upstream influence of Tau in the cancer biology of P53, an influence that may lose its weight when P53 is mutated. To investigate this aspect, we evaluated *MAPT* expression in P53 wild-type (WT) versus mutated tumors (**Figure 2a**). The expression of *MAPT* was significantly different in WT and P53-mutated tumors in four cancer types. We observed a highly significant decrease in *MAPT* expression in P53-mutated BRCA, whereas there was a significant increase in *MAPT* expression in P53-mutated LGG, PAAD, and LIHC. In BRCA it has been shown that the expression of *MAPT* is associated with subtypes i.e., high in ER+ tumors which have the lowest P53 mutation rate(40). P53 mutations were additionally functionally grouped in truncating mutation/homozygous deletion (Truncating/HomDel) and inframe/missense mutations, which could have a distinct functional impact and, in this case, a distinct impact on *MAPT*. No major differences were identified, except for SARC and OV (**Figure S4**). In SARC there is a lower expression of *MAPT* in tumors with Truncating/HomDel mutations than in tumors with in-frame/missense mutations, whereas in OV the opposite behaviour is detected.

**Figure S4.**
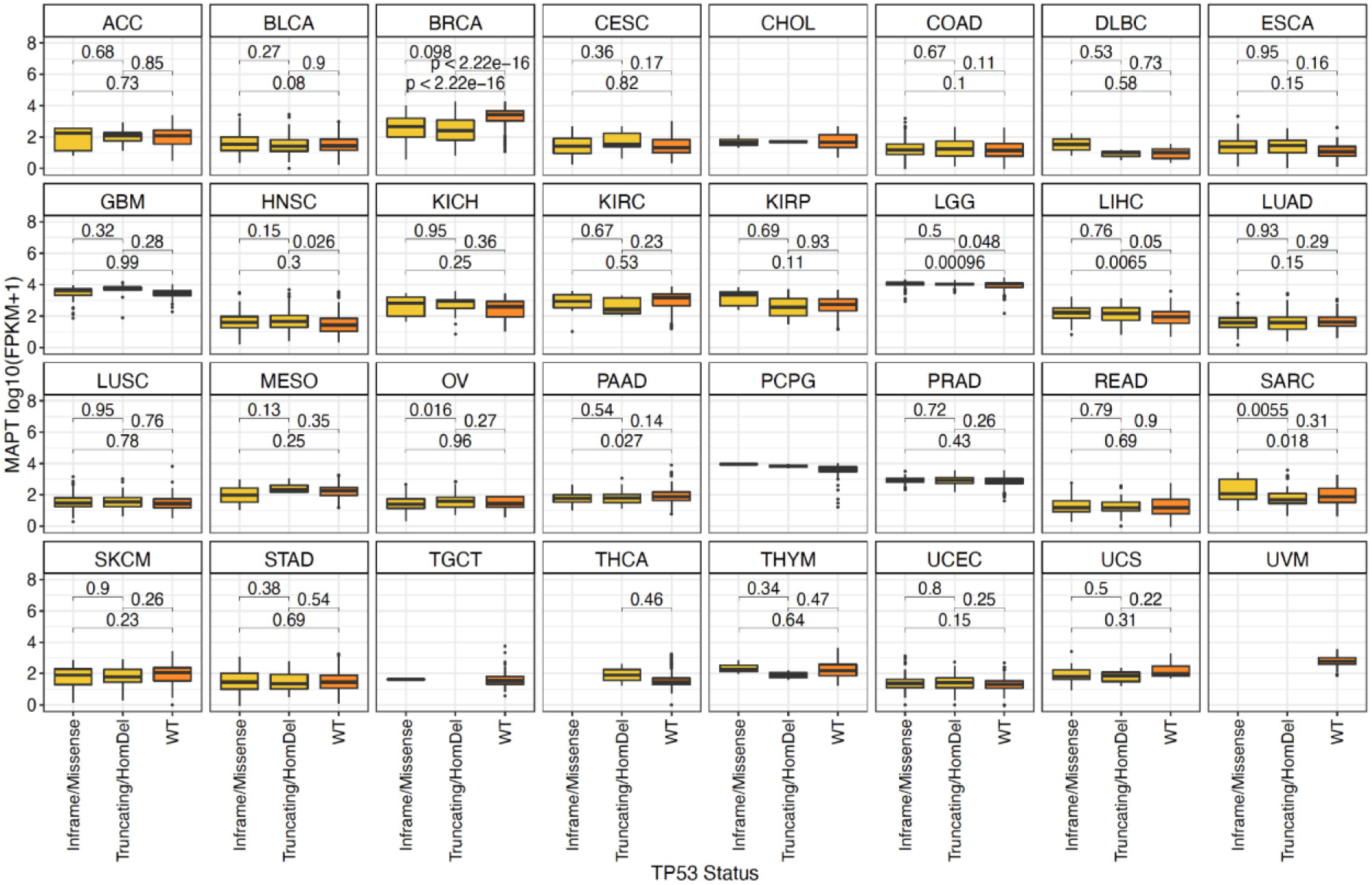
**-** Expression of MAPT in tumors with WT P53 or with either a Truncating/HomDel mutation or an Inframe/Missense mutation in PT53. Differences were evaluated by a two-sided Student’s t-test.

**Table S1.**
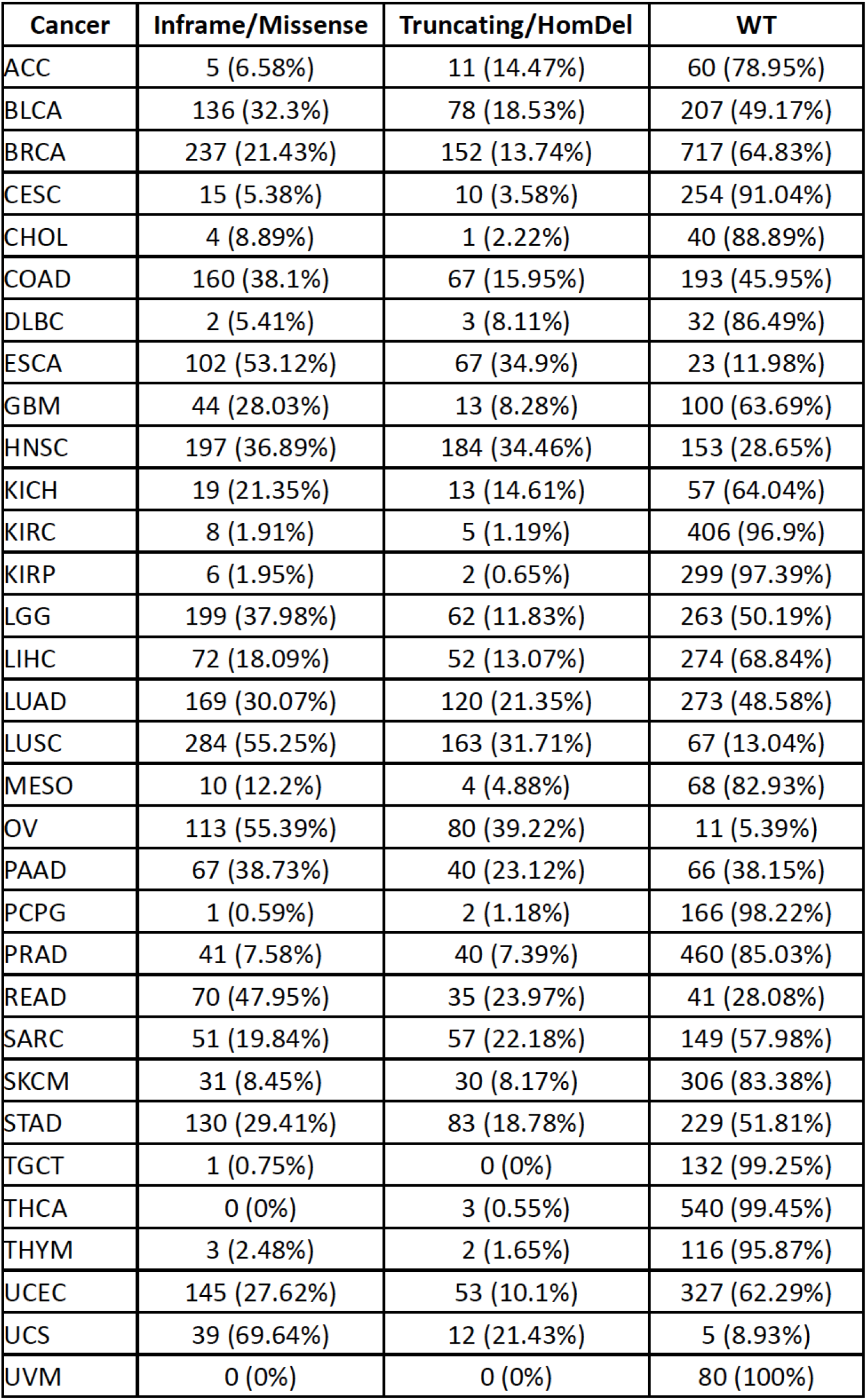
**-** Percentage of TP53 mutations per cancer type. Mutation frequency is separated according to the different mutation types (inframe/missense vs truncating/homdel).

| Cancer | Inframe/Missense | Truncating/HomDel | WT |
| --- | --- | --- | --- |
| ACC | 5 (6.58%) | 11 (14.47%) | 60 (78.95%) |
| BLCA | 136 (32.3%) | 78 (18.53%) | 207 (49.17%) |
| BRCA | 237 (21.43%) | 152 (13.74%) | 717 (64.83%) |
| CESC | 15 (5.38%) | 10 (3.58%) | 254 (91.04%) |
| CHOL | 4 (8.89%) | 1 (2.22%) | 40 (88.89%) |
| COAD | 160 (38.1%) | 67 (15.95%) | 193 (45.95%) |
| DLBC | 2 (5.41%) | 3 (8.11%) | 32 (86.49%) |
| ESCA | 102 (53.12%) | 67 (34.9%) | 23 (11.98%) |
| GBM | 44 (28.03%) | 13 (8.28%) | 100 (63.69%) |
| HNSC | 197 (36.89%) | 184 (34.46%) | 153 (28.65%) |
| KICH | 19 (21.35%) | 13 (14.61%) | 57 (64.04%) |
| KIRC | 8 (1.91%) | 5 (1.19%) | 406 (96.9%) |
| KIRP | 6 (1.95%) | 2 (0.65%) | 299 (97.39%) |
| LGG | 199 (37.98%) | 62 (11.83%) | 263 (50.19%) |
| LIHC | 72 (18.09%) | 52 (13.07%) | 274 (68.84%) |
| LUAD | 169 (30.07%) | 120 (21.35%) | 273 (48.58%) |
| LUSC | 284 (55.25%) | 163 (31.71%) | 67 (13.04%) |
| MESO | 10 (12.2%) | 4 (4.88%) | 68 (82.93%) |
| OV | 113 (55.39%) | 80 (39.22%) | 11 (5.39%) |
| PAAD | 67 (38.73%) | 40 (23.12%) | 66 (38.15%) |
| PCPG | 1 (0.59%) | 2 (1.18%) | 166 (98.22%) |
| PRAD | 41 (7.58%) | 40 (7.39%) | 460 (85.03%) |
| READ | 70 (47.95%) | 35 (23.97%) | 41 (28.08%) |
| SARC | 51 (19.84%) | 57 (22.18%) | 149 (57.98%) |
| SKCM | 31 (8.45%) | 30 (8.17%) | 306 (83.38%) |
| STAD | 130 (29.41%) | 83 (18.78%) | 229 (51.81%) |
| TGCT | 1 (0.75%) | 0 (0%) | 132 (99.25%) |
| THCA | 0 (0%) | 3 (0.55%) | 540 (99.45%) |
| THYM | 3 (2.48%) | 2 (1.65%) | 116 (95.87%) |
| UCEC | 145 (27.62%) | 53 (10.1%) | 327 (62.29%) |
| UCS | 39 (69.64%) | 12 (21.43%) | 5 (8.93%) |
| UVM | 0 (0%) | 0 (0%) | 80 (100%) |

**Figure 2.**
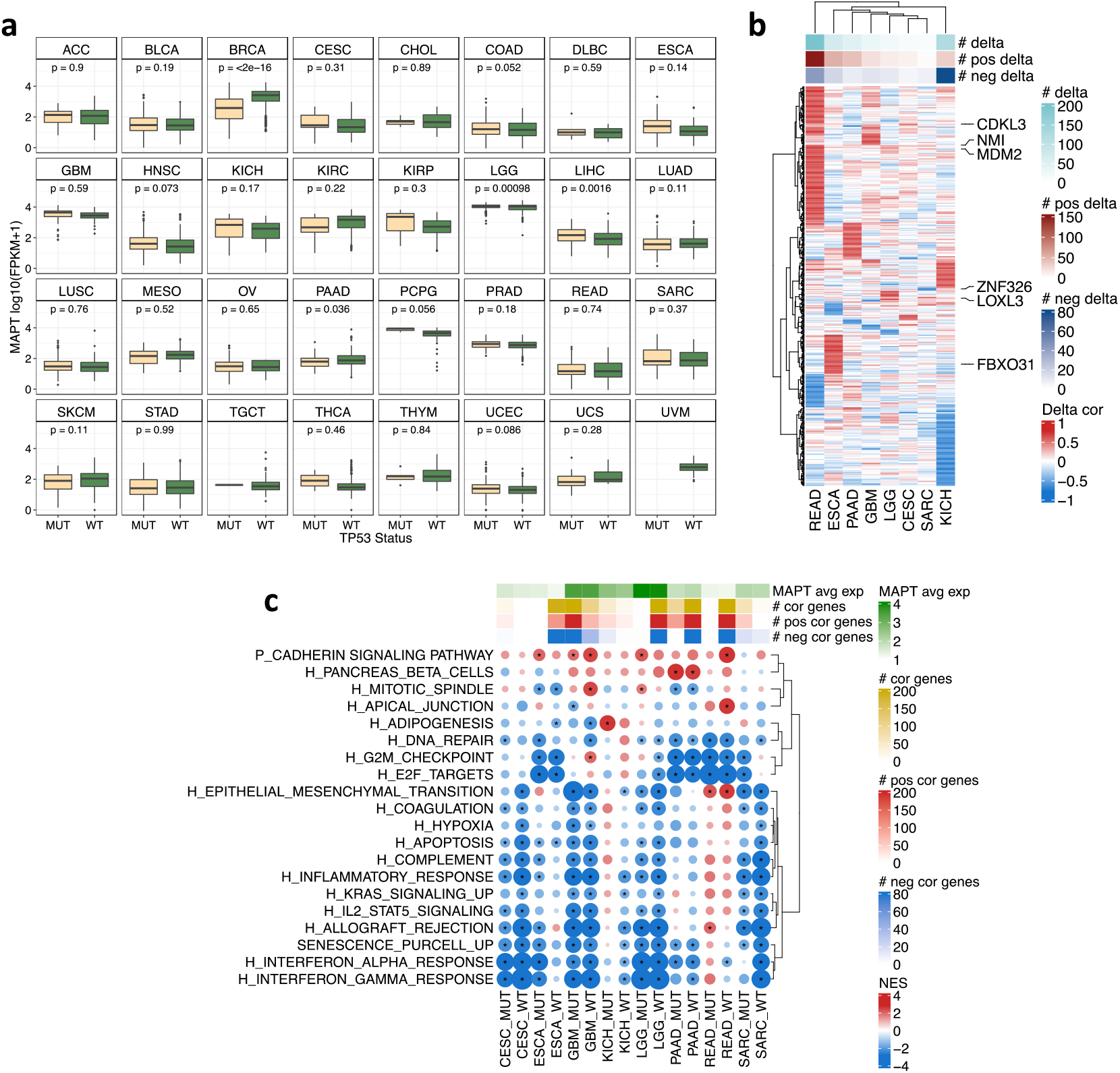
- Pan-cancer evaluation of MAPT expression and transcriptional associations stratified by P53 status. **(a)** Expression of MAPT in TP53 WT and TP53 MUT tumors. The analysis is stratified by cancer type. Differences are evaluated by a two-sided Student’s t-test. **(b)** Heatmap of 514 genes with an absolute delta correlation above 0.6 in P53 WT vs P53 MUT tumors for at least one cancer type. The correlation analysis was performed only for cancer types that included > 20 patients for each P53 status. Eight cancer types had more than one gene with a significant delta and are shown. An extended version of the results is reported in Figures S5 and S6. **(c)** GeneSet Enrichment Analysis on the genes ranked accordingly to their correlation with Tau expression within each cancer type and separately for P53 wild-type and P53-mutant tumors. Genesets with an absolute delta enrichment above 2.3 are reported for the eight cancer types shown in (b).

Next, we separately delineated genes and pathways associated with *MAPT* expression in 19 cancer types with at data from at least 20 tumors with mutated or WT P53 (**Table S1**). We subsequently searched for genes correlated with *MAPT* expression, separately for P53-mutated and WT tumors, and computed, for each cancer type, their correlation delta (**Figure S5**). Significant delta values (i.e. absolute delta above 0.6) were found in 8 cancer types (READ, ESCA, PAAD, GBM, LGG, CESC, SARC, and KICH) and 514 genes had a significant delta in at least one cancer type (**Figure 2b**, **Figure S6**, and **Figure S7**). For three cancer types, i.e. CESC, ESCA, and READ, while no genes were significantly correlated with *MAPT* in the overall population (**Figure S2**), correlated genes emerged when stratifying for P53 status. In general, we observed a prevalence of positive delta (i.e. higher correlation with *MAPT* in P53 mutant). This was particularly evident in READ, while a prevalence of negative delta was observed in KICH (**Figure 2b**).

**Figure S5.**
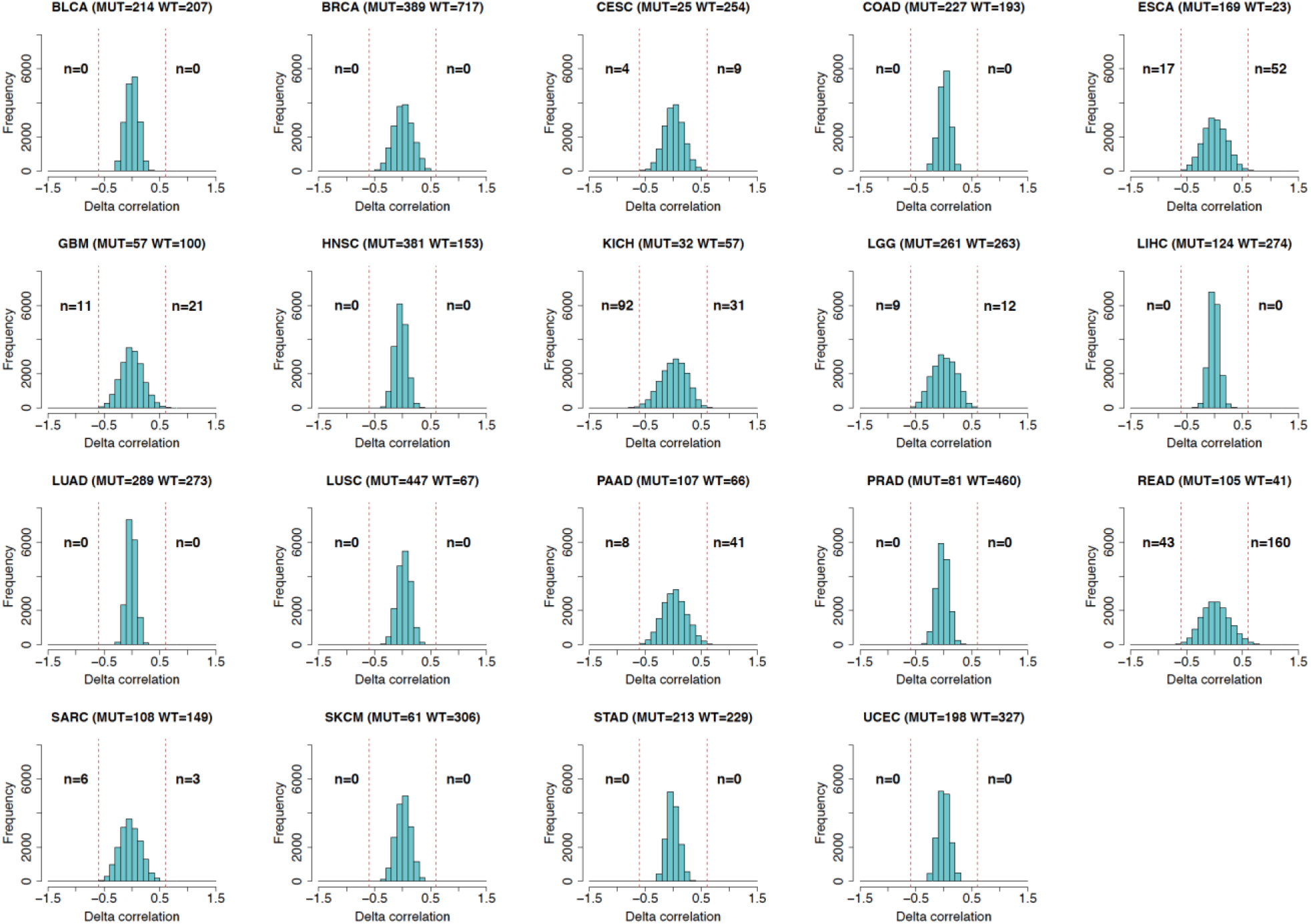
**-** Correlation with MAPT for all genes was computed separately for TP53 MUT and TP53 WT tumors. The delta correlation distribution (correlation in TP53 MUT – correlation in TP53 WT) is reported in the figure for each cancer type. The number of genes with an absolute delta above 0.6 is indicated.

This analysis allowed us to highlight *MAPT*-gene correlations that strongly depended on the P53 status, identifying an additional pool of genes with a potential context-specific biological link with *MAPT*. Among them, MDM2, a key P53 antagonist, displayed a positive delta in seven out of eight cancers. The correlation between *MAPT* and *MDM2* was mostly negative in WT P53 tumors whereas mutant P53 tumors showed either a loss of correlation (GBM, LGG) or a correlation in the opposite way (READ, CESC, SARC). Since a loss of correlation with mutated P53 may indicate a role for Tau upstream of P53, and considering that MDM2 is the main E3 protein ubiquitin ligase and inhibitor of P53, the Tau-MDM2 link may suggest a possible mechanism for the modulatory role of Tau on P53(16). Alternatively, considering that P53 activation of *MDM2* transcription(41) is lost when P53 is mutated, this may explain the reduced *MAPT*-*MDM2* correlation when P53 is mutated in LGG and GBM.

A single gene: *NMI* (N-Myc and STAT interactor), showed a weak positive delta in all eight cancer types with a negative correlation between *MAPT* and *NMI* in WT P53 tumors in all eight cancer types and a decrease of the correlation in P53 mutant tumors. NMI is an interferon-inducible protein participating in various cellular activities and has been involved in the process of tumorigenesis and tumor progression(42). The associations of *MAPT* expression with genes encoding for proteins involved in proliferation, EMT, and inflammation were very variable among cancers. As an example, the correlations with *CDKL3* and *FBXO31*, two genes encoding for proteins involved in proliferation show similar behavior in READ and ESCA, with a negative correlation in WT P53 tumors and a positive correlation in P53 mutant tumors, whereas quite opposite correlation changes were observed in GBM, CESC and KICH. In the EMT dataset, some correlations with *MAPT* vary similarly in a defined cancer type whereas they change oppositely in another cancer type. As an example, a negative correlation is detected with *LOXL3* and *ZNF326* in P53 WT tumors and these correlations are strongly decreased in mutant P53 tumors, whereas in SARC, *MAPT* negatively correlates with *LOXL3* and positively with *ZNF326*, and these correlations are reversed in mutant P53 tumors.

In the eight cancer types with significant delta values, we complemented the gene level analysis with a GSEA on genes ranked according to the correlation with *MAPT* in P53 WT and P53 mutated tumors for the same eight cancers. Genesets with a significant delta NES are shown in **Figure 2c**. Several enrichments are affected by P53 status, positively or negatively, and in a cancer-specific manner. For CESC, ESCA, GBM, and LGG, the association of *MAPT* with inflammatory pathways is not changed according to P53 status. On the contrary, IFN-related genesets had a negative association with *MAPT* expression in KICH and SARC overall population (**Figure 1d**); however, after stratifying by P53 status, we found that such association was stronger and significant only in P53 WT tumors. Also, the association of cell cycle genesets with *MAPT* was affected by P53 status in some cancer types. Indeed, the positive association with *MAPT* in GBM (G2M_CHECKPOINT) was limited to the P53 WT tumors; similarly, the negative association in LGG was limited to P53 WT tumors. In SARC, no association with cell cycle genesets was observed overall (**Figure 1d**); however, a significant negative association was present in P53 mutant tumors.

These data indicated that *MAPT*-correlated modulation of several biological processes depended on the status of P53 for some cancer types, possibly informative for an upstream or downstream effect of Tau on P53 cancer biology.

**Figure S6.**
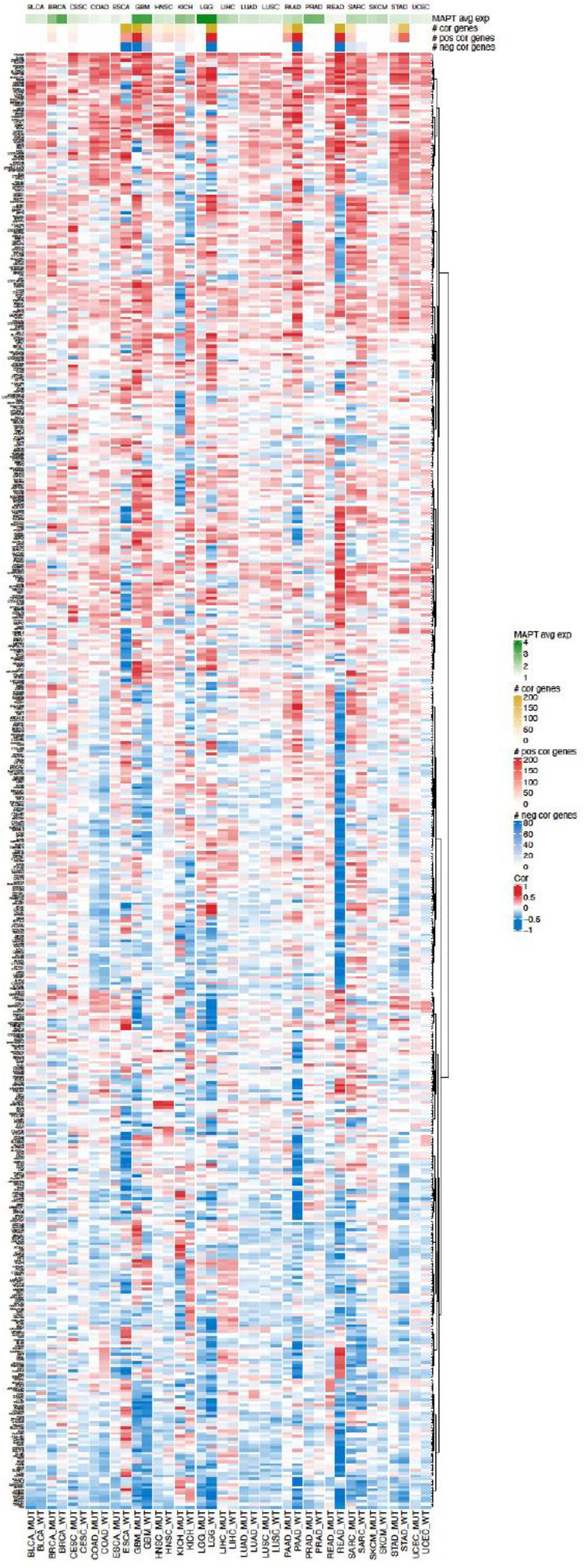
**-** Heatmap of 514 genes with an absolute delta correlation above 0.6 in P53 WT vs P53 MUT tumors for at least one cancer type. The correlation analysis was performed only for 19 cancer types with at least 20 patients with P53 WT and 20 P53 MUT. MAPT correlation values are shown. For each group identified by the combination of cancer type and P53 status, the average MAPT expression and the number of genes correlated with MAPT are shown.

**Figure S7.**
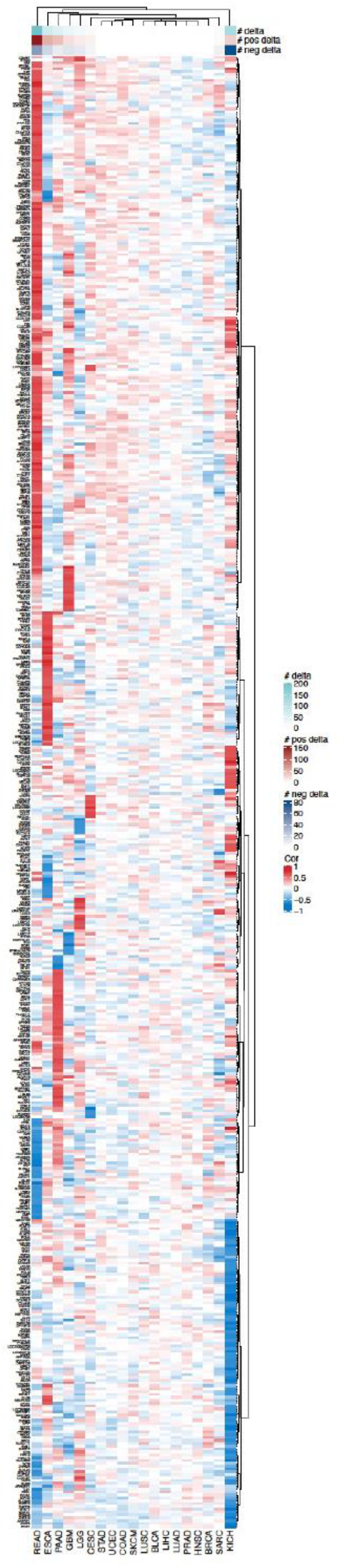
**-** Heatmap of 514 genes with an absolute delta correlation above 0.6 in P53 WT vs P53 MUT tumors for at least one cancer type. The correlation analysis was performed for 19 cancer types with at least 20 patients with P53 WT and 20 P53 MUT.

### Cancer-specific and P53-dependent associations of *MAPT* with patients’ survival

After exploring genes and pathways linked to *MAPT* expression, we thoroughly analyzed the association of *MAPT* with cancer survival. Univariate Cox regression analysis was performed for each cancer in the overall population, and separately for P53 WT and mutant subgroups. Then, we applied a multivariate Cox regression analysis adjusting for tumor size, lymph node status, metastatic status, and expression of the *AURKA* gene as a reporter of the proliferation process, associated with survival in multiple cancers (**Table 1**, **Figure S9**).

**Table 1.** **-** Univariate and multivariate Cox survival analysis for each cancer type in all samples and stratified by P53 status.

| Cancer type | ALL - Univariate |  |  | ALL - Multivariate |  |  | TP53 WT - Univariate |  |  | TP53 WT - Multivariate |  |  | TP53 MUT - Univariate |  |  | TP53 MUT - Multivariate |  |  |
| --- | --- | --- | --- | --- | --- | --- | --- | --- | --- | --- | --- | --- | --- | --- | --- | --- | --- | --- |
|  | N | HR | P | N | HR | P | N | HR | P | N | HR | P | N | HR | P | N | HR | P |
| ACC | 79 | 0.97 | 0.9418 | 77 | 0.76 | 0.5226 |  |  |  |  |  |  |  |  |  |  |  |  |
| BLCA | 423 | 1.22 | 0.1753 | 179 | 1.59 | 0.0516 | 202 | 1.33 | 0.1903 | 95 | 2.09 | 0.0475 | 215 | 1.12 | 0.5861 | 80 | 1.28 | 0.4729 |
| BRCA | 1203 | 0.30 | 7.96E-09 | 1017 | 0.31 | 8.10E-08 | 704 | 0.28 | 3.49E-05 | 585 | 0.31 | 0.0006 | 387 | 1.32 | 0.3093 | 344 | 0.88 | 0.7065 |
| CESC | 296 | 1.30 | 0.2976 | 104 | 1.06 | 0.9208 | 241 | 1.19 | 0.5324 | 81 | 0.93 | 0.9244 |  |  |  |  |  |  |
| CHOL | 45 | 0.80 | 0.6037 | 38 | 0.97 | 0.9553 |  |  |  |  |  |  |  |  |  |  |  |  |
| COAD | 470 | 1.64 | 0.0161 | 408 | 1.20 | 0.4606 | 178 | 1.46 | 0.2915 | 156 | 0.69 | 0.4384 | 221 | 2.10 | 0.0100 | 183 | 1.59 | 0.1692 |
| ESCA | 196 | 1.04 | 0.8781 | 149 | 0.88 | 0.6761 |  |  |  |  |  |  | 169 | 1.13 | 0.6239 | 128 | 0.92 | 0.7954 |
| GBM | 167 | 0.97 | 0.8589 | 167 | 0.98 | 0.9053 | 100 | 1.04 | 0.8620 | 100 | 1.04 | 0.8558 | 57 | 1.05 | 0.8723 | 57 | 1.10 | 0.7623 |
| HNSC | 562 | 1.43 | 0.0080 | 182 | 1.68 | 0.0511 | 153 | 1.24 | 0.4567 | 105 | 1.57 | 0.2252 | 378 | 1.33 | 0.0666 | 133 | 1.49 | 0.2048 |
| KICH | 89 | 1.50 | 0.5212 | 37 | 1.19 | 0.9264 |  |  |  |  |  |  |  |  |  |  |  |  |
| KIRC | 604 | 0.44 | 3.94E-07 | 293 | 0.61 | 0.0598 |  |  |  |  |  |  |  |  |  |  |  |  |
| KIRP | 320 | 1.26 | 0.4477 | 59 | 2.44 | 0.1254 |  |  |  |  |  |  |  |  |  |  |  |  |
| LGG | 529 | 0.24 | 2.49E-10 | 529 | 0.34 | 4.87E-06 | 260 | 0.19 | 3.94E-07 | 260 | 0.33 | 0.0018 | 260 | 0.29 | 0.0002 | 260 | 0.36 | 0.0030 |
| LIHC | 418 | 1.46 | 0.0209 | 262 | 1.22 | 0.3672 | 269 | 1.15 | 0.5209 | 172 | 1.01 | 0.9699 | 123 | 1.69 | 0.0712 | 91 | 1.37 | 0.3972 |
| LUAD | 563 | 0.77 | 0.0734 | 393 | 0.90 | 0.5329 | 265 | 0.67 | 0.0653 | 190 | 0.87 | 0.5777 | 284 | 0.87 | 0.4713 | 194 | 0.98 | 0.9510 |
| LUSC | 548 | 1.17 | 0.2538 | 443 | 1.07 | 0.6559 | 66 | 1.23 | 0.5628 | 65 | 1.43 | 0.3424 | 443 | 1.18 | 0.3084 | 362 | 1.15 | 0.4394 |
| MESO | 86 | 1.17 | 0.5212 | 57 | 1.05 | 0.8766 |  |  |  |  |  |  |  |  |  |  |  |  |
| OV | 308 | 1.19 | 0.2809 | 308 | 1.19 | 0.2751 |  |  |  |  |  |  |  |  |  |  |  |  |
| PAAD | 182 | 0.68 | 0.0663 | 85 | 1.10 | 0.7659 | 65 | 0.61 | 0.2075 | 61 | 0.68 | 0.4290 | 107 | 0.71 | 0.1802 | 53 | 0.92 | 0.8283 |
| READ | 164 | 1.00 | 0.9918 | 147 | 0.87 | 0.7511 |  |  |  |  |  |  | 100 | 2.87 | 0.1105 | 91 | 1.99 | 0.3267 |
| SARC | 265 | 1.10 | 0.6614 | 265 | 1.22 | 0.3725 | 148 | 1.05 | 0.8579 | 148 | 1.07 | 0.8185 | 109 | 1.16 | 0.6691 | 109 | 1.55 | 0.2657 |
| SKCM | 463 | 1.03 | 0.8556 | 357 | 1.35 | 0.1224 | 293 | 1.43 | 0.0780 | 219 | 2.06 | 0.0035 | 64 | 0.98 | 0.9554 | 51 | 1.16 | 0.7630 |
| STAD | 421 | 1.12 | 0.4535 | 392 | 1.01 | 0.9357 | 208 | 1.09 | 0.6750 | 194 | 0.96 | 0.8735 | 208 | 1.07 | 0.7609 | 193 | 1.01 | 0.9563 |
| THCA | 571 | 2.11 | 0.1356 | 324 | 3.57 | 0.1182 |  |  |  |  |  |  |  |  |  |  |  |  |
| UCEC | 553 | 1.94 | 0.0041 | 553 | 2.01 | 0.0026 | 325 | 2.59 | 0.0161 | 325 | 2.80 | 0.0093 | 198 | 1.87 | 0.0415 | 198 | 1.84 | 0.0471 |
| UCS | 56 | 1.87 | 0.0830 | 56 | 1.84 | 0.0925 |  |  |  |  |  |  |  |  |  |  |  |  |
| UVM | 80 | 2.66 | 0.1031 | 55 | 0.27 | 0.1568 |  |  |  |  |  |  |  |  |  |  |  |  |

A high expression of *MAPT* was associated with a worse prognosis in COAD, HNSC, LIHC, and UCEC in the univariate analysis, and only for UCEC in the multivariate analysis. In contrast, high *MAPT* expression was associated with a better prognosis in BRCA, KIRC, and LGG in the univariate analysis and for BRCA and LGG in the multivariate analysis. These data suggested that the (positive or negative) correlation of *MAPT* expression with survival was independent of the other prognostic factors for BRCA, LGG, and UCEC. For some examples, we found a P53 status-dependent association between *MAPT* expression and cancer survival. This was the case for BRCA where the *MAPT*-survival correlation was lost for the P53 mutant cases. In other cancer types, for example, LGG, the positive association of *MAPT* with survival occurred in both P53 WT and P53 mutant cohorts. Similarly, for UCEC, the negative association appeared independent of the P53 status. Finally, the negative association between *MAPT* and survival detected for COAD appeared limited to the P53 mutant cases. The most significant Kaplan-Meier curves showing the association between *MAPT* and survival are shown in **Figure 3a** and the complete analysis is reported in **Figure S8**.

**Figure 3.**
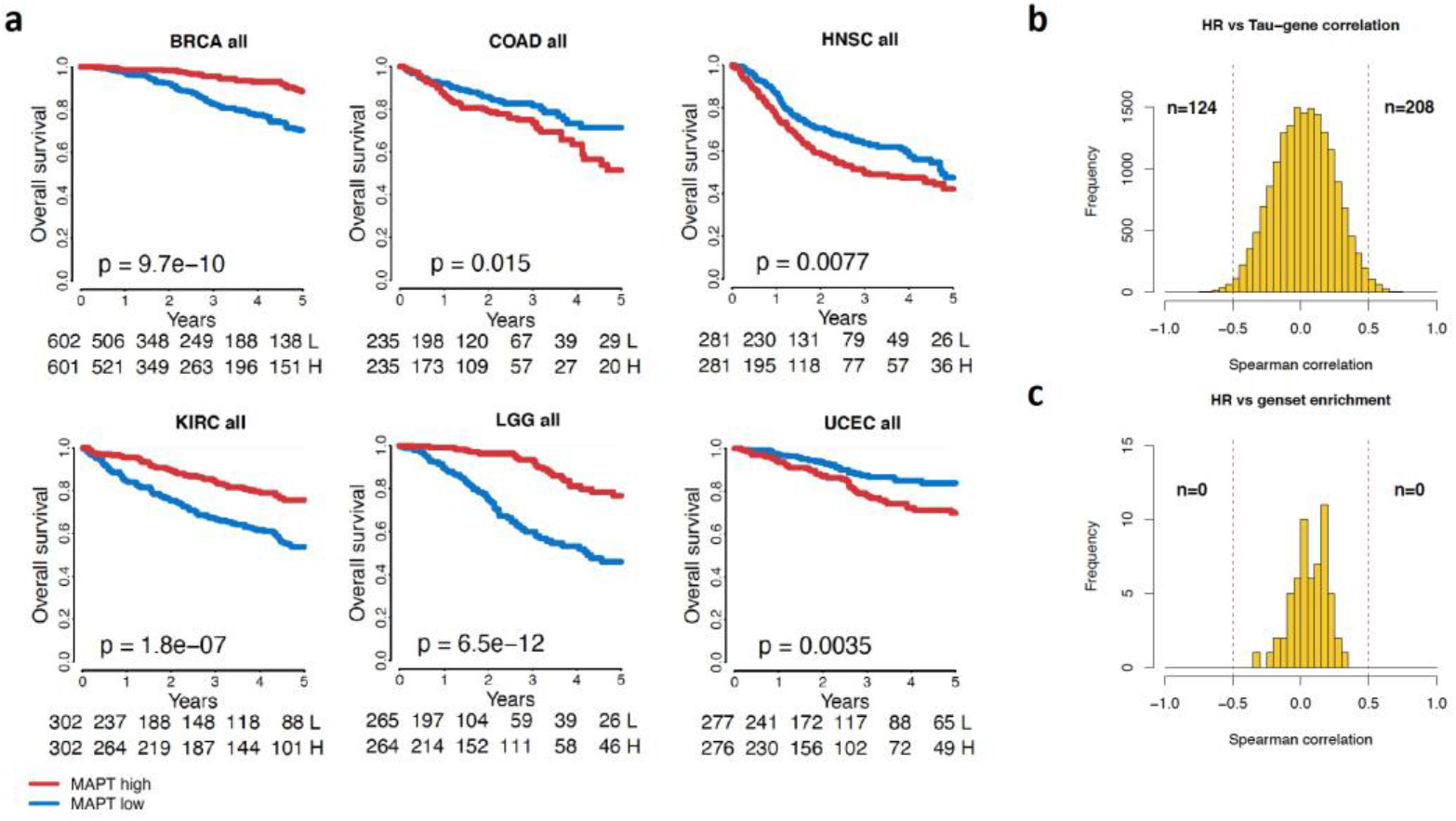
- MAPT cancer-specific association with survival. **(a)** Selected Kaplan-Meier curves showing the association between MAPT and survival in 6 cancer types. P-value obtained by log-rank test. **(b)** For each gene, its correlation with MAPT across all cancer types was correlated with hazard ratios (univariate Cox analysis) for the same cancer types. Histogram shows the distribution of values obtained. Number of genes with a correlation > 0.5 or < -0.5 are indicated. **(c)** Same analysis as in (b) but computed correlating NES for each geneset with hazard ratios.

Since the variable association with survival in different cancer types, we evaluated whether this could be related to a change in the biological network associated with *MAPT* expression. To this aim, we searched for a correlation between *MAPT*-gene correlation values and *MAPT* hazard ratios from the Cox univariate analysis. We found associations between the way *MAPT* correlated with certain genes and the way it was associated with survival (**Figure 3b**). Among the top genes, some encode for fundamental structural proteins such as a collagen subunit (*COL5A3*) or an extracellular matrix glycoprotein (*THSD4*), which play a weighty role in EMT and migration processes(43, 44). Other genes (e.g., *FBXL16*, *PDIA6*, *PPIB*) are involved in processes of protein folding and/or degradation, and their possible implication in tumorigenesis has been reported (45–47) (**Figure S10**). A similar analysis was done to link hazard ratios with GSEA enrichment scores but no geneset was identified (**Figure 3c**).

**Figure S8.**
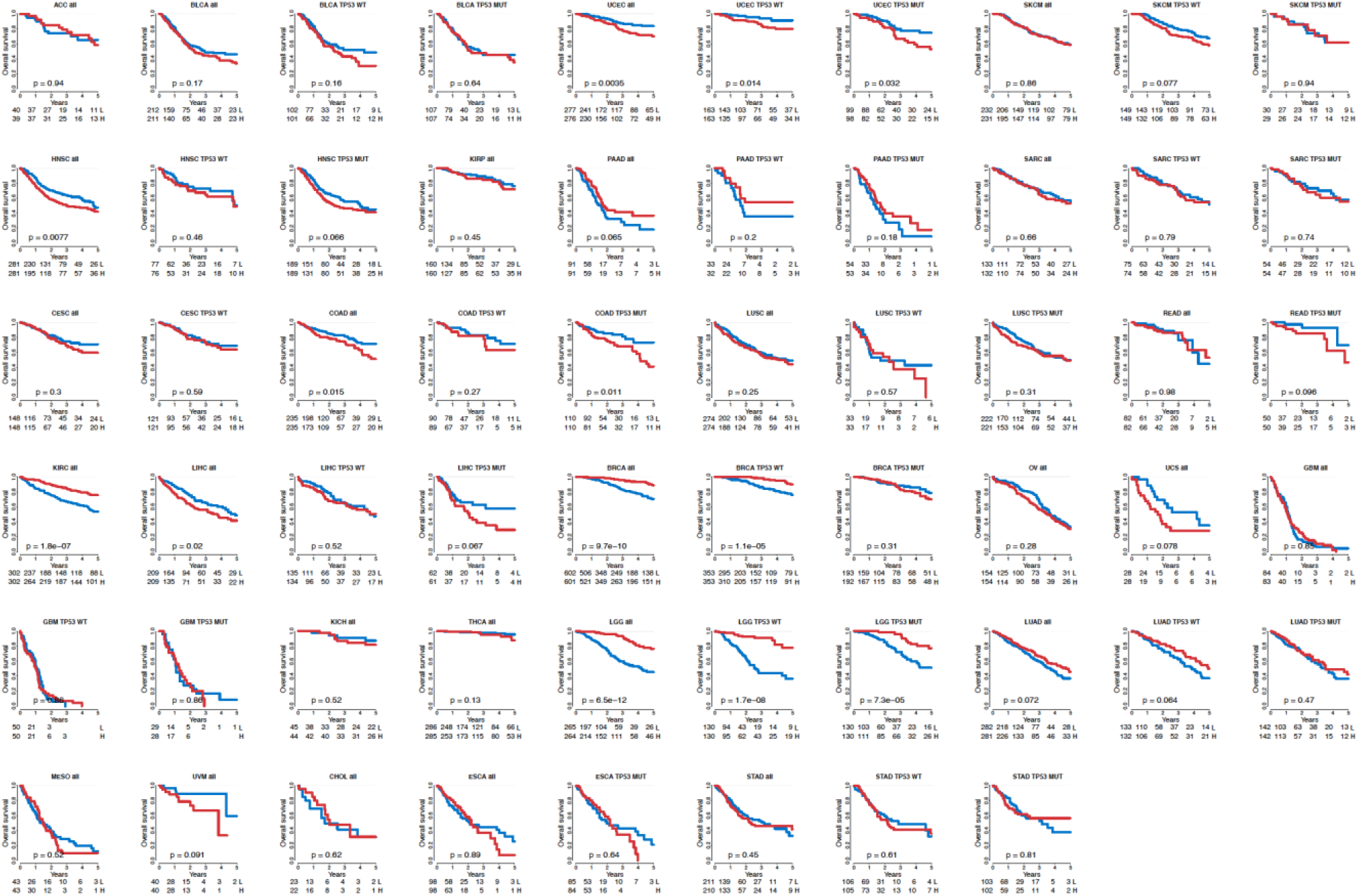
- MAPT cancer-specific association with survival. Kaplan-Meier curves showing the association between MAPT (High/Low using the median as threshold) and survival in the overall population and stratifying according to P53 status. P-value obtained by log-rank test.

**Figure S9.**
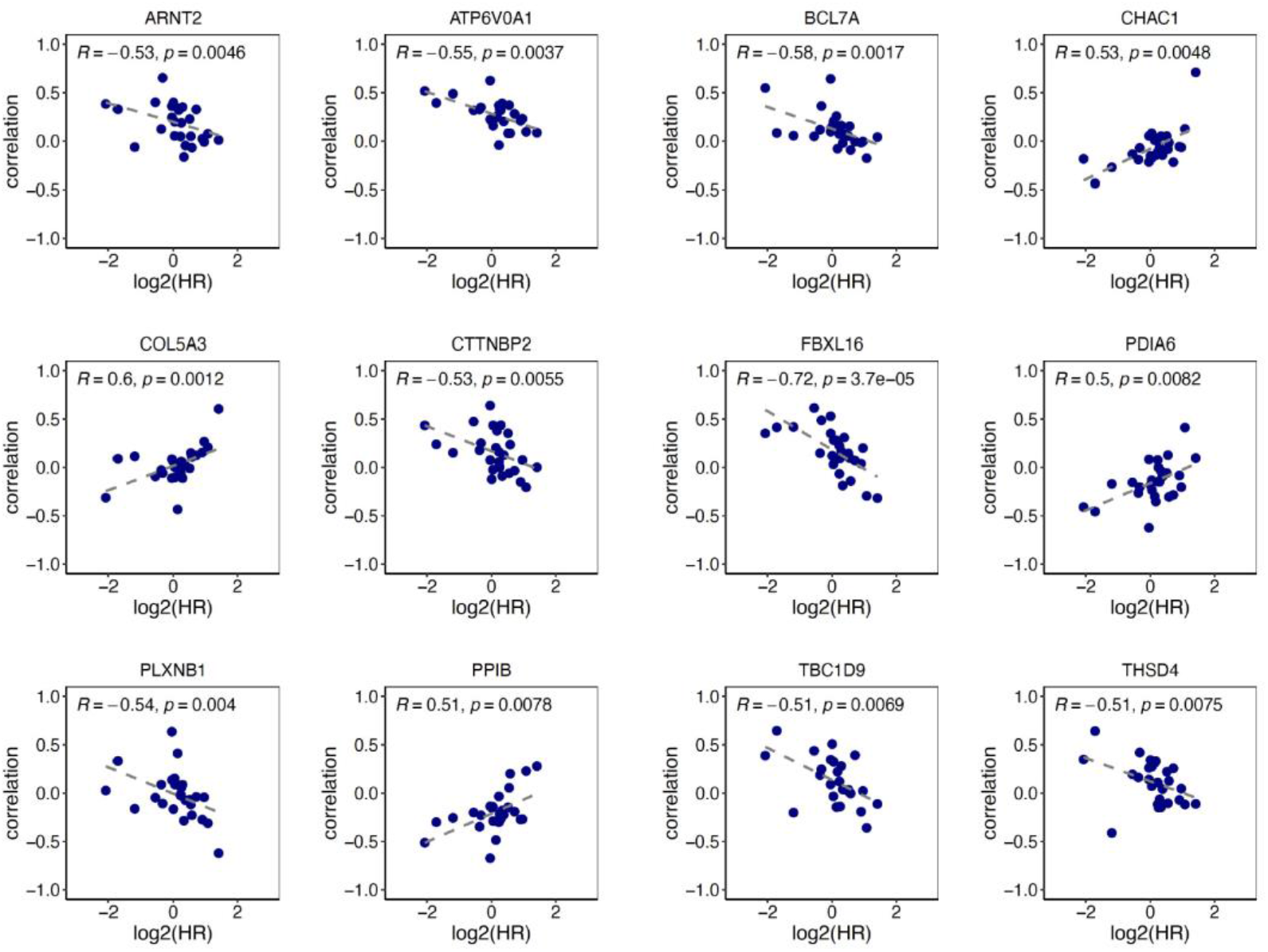
**-** Scatterplots for genes showing an absolute correlation >0.5 between the MAPT-gene correlation and MAPT hazard ratio across different cancer types. The 12 genes achieving an absolute MAPT-gene correlation >0.6 (as in **Figure 1c**) are shown.

### *MAPT* expression, essentiality, and association with drug response in pre-clinical models

Our analysis to chart the relevance of *MAPT* in cancer continued in pan-cancer pre-clinical data. We analyzed the CRISPR DEPMAP data collection (https://depmap.org/portal/) for the cell viability after *MAPT* KO across cell lines from 29 different cancer types. The impact of *MAPT* KO on viability was cancer-specific but overall, a decrease in viability is prevalently observed, except for rhabdoid cell lines **(Figure 4a)**.

**Figure 4.**
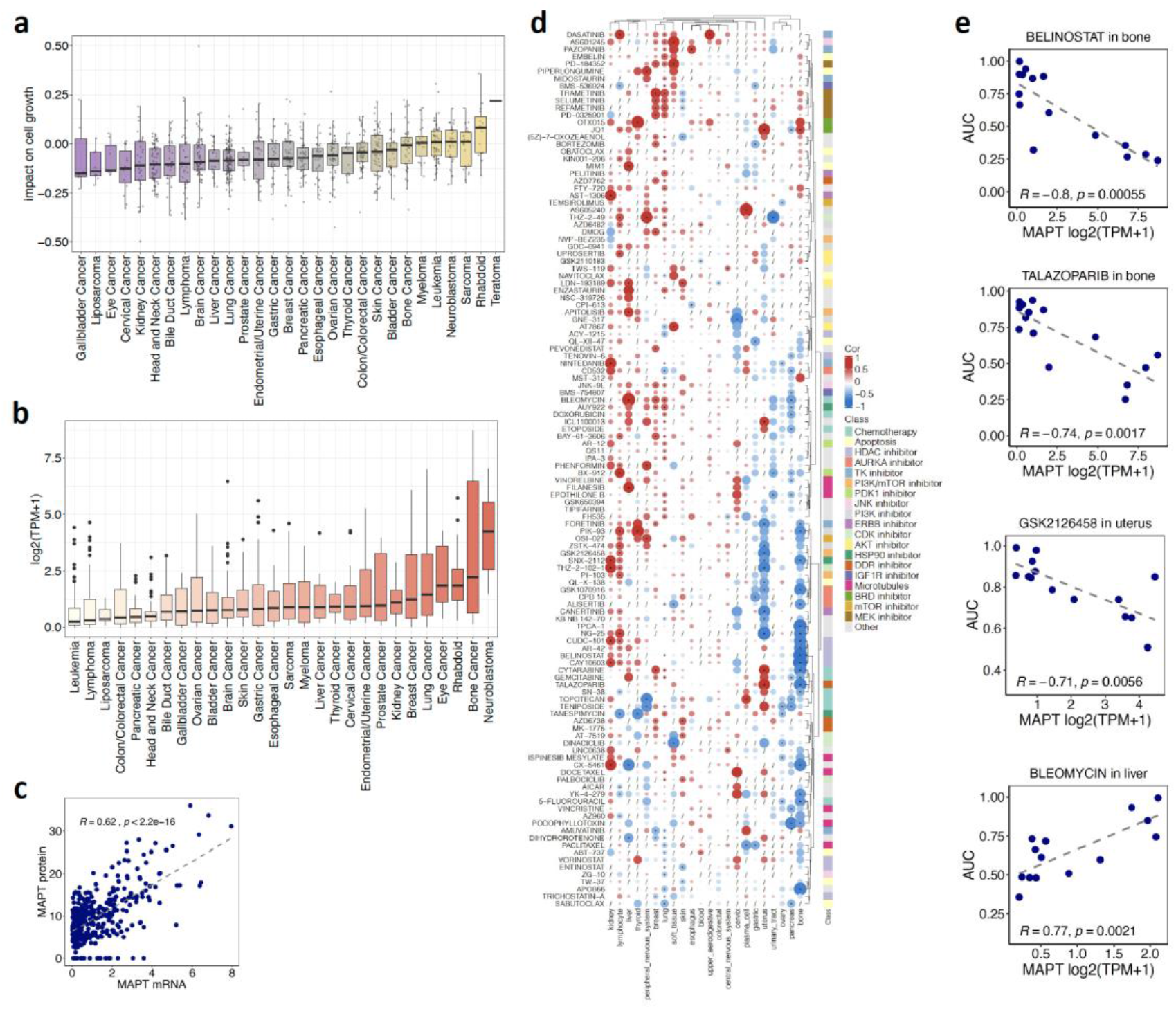
- MAPT expression, essentiality, and association with drug response in pre-clinical models. **(a)** Cell viability after MAPT KO in the CRISPR DEPMAP dataset. Viability scores are normalized such that nonessential genes have a median score of 0 and independently identified common essentials have a median score of -1. **(b)** MAPT expression in the DEPMAP cell line dataset according to the cancer type. **(c)** Correlation of MAPT gene expression with Tau protein (sum of three isoforms) in the DEPMAP cell line dataset. **(d)** Heatmap summarizing the correlations between MAPT expression with drug response quantified as area under the drug response curve (AUC). **(e)** Selected scatterplots of MAPT-drug response association.

For the DEPMAP cell line data collection, we also correlated *MAPT* expression with drug response, where the area under the drug response curve (AUC) with values ∼0 indicated drug sensitivity and values ∼1 indicated drug resistance. First, we evaluated *MAPT* expression across cancer cell lines grouped by cancer type. The highest expression was observed in neuroblastoma and bone cancer (**Figure 4b**). The availability of proteomic data in the DEPMAP dataset for a subset of cell lines allowed us to determine that *MAPT* transcript and Tau protein strongly correlated (cor = 0.62), indicating that evaluating mRNA expression is informative for Tau protein levels (**Figure 4c**).

After data filtering, we evaluated 121 drugs in 22 cancers (**Figure 4d-e**). The associations involved multiple drug families and were largely specific to the cancer type. Of note, drugs tended to cluster based on their target or mechanism. A negative correlation between *MAPT* expression and resistance to several drugs with various modes of action is detected in pancreatic, uterine, and mainly bone cancer-derived cell lines, whereas predominantly positive correlations were found in cell lines derived from other cancers. In bone cancer cell lines, a high *MAPT* expression was associated with response to kinase inhibitors (Aurora, PI3K), HDAC inhibitors as well as DNA damaging drugs. The most significant correlations were found with belinostat, an HDAC inhibitor mainly used for the treatment of peripheral T cell lymphoma(48), and with talazoparib, a PARP inhibitor used for the treatment of advanced breast cancer with germline BRCA mutation(49). Both drugs are now in clinical studies in bone cancers(50, 51). In liver cancer cell lines, *MAPT* expression was negatively associated with resistance to bleomycin, a DNA-damaging drug approved for squamous cell head and neck cancer, Hodgkin’s lymphoma, and testicular carcinoma(52). In uterine cancer cells, high *MAPT* expression correlated with sensitivity to kinase inhibitors. A very good correlation resulted with GSK2126458, a PI3K/mTOR inhibitor with broad antitumor activity in preclinical and clinical studies(53). Resistance to MEK inhibitors correlated positively with *MAPT* for cell lines derived from breast and lung cancer, and negatively for skin-derived cell lines. Association between *MAPT* expression and resistance to kinases inhibitors was also detected in lymphocytes (PI3K) and bone and uterus-derived cancer cell lines (Aurora kinase). Surprisingly, response to microtubule-targeting drugs did not show a very strong correlation with *MAPT* expression. While cells derived from uterine and bone cancers were the ones with the higher number of drugs correlated with *MAPT* expression, only a few drugs did the same for cell lines derived from tumors affecting tissues such as the CNS, colorectal, upper aerodigestive, blood, and esophagus. Cell lines from the cervix, soft tissue, lung, breast, peripheral nervous system, liver, lymphocyte, and kidney showed a *MAPT* correlation with the response to several drugs.

Overall, these data highlighted that *MAPT* expression may represent an informative predictor of drug response for multiple cancer types.

## Discussion

To strengthen the emerging evidence for the role of *MAPT* in cancer, we performed a pan-cancer *in silico* analysis to define the landscape of pathways, genes, and drug treatments associated with *MAPT* expression. We report a highly significant association between *MAPT* and cell proliferation, inflammation, and EMT-related genes. These genes are part of cellular pathways fundamental for tumor initiation, progression, and heterogeneity(54), with the latter also having a key link with the tumor microenvironment. In particular, interferons and inflammation-related genes could suggest a link between *MAPT* expression and inhibition of anti-tumor immune response, with possible therapeutic implications given the fast-paced clinical implementation of the immunotherapy (55).

Interestingly, we observed a positive pan-cancer correlation of *MAPT* with a large set of neuronal genes. Some of these have a clear role in AD pathology(35), further connecting these two very different human conditions that are usually considered to be inversely associated(56). These findings are also in line with evidence suggesting that cancer cells recapitulate features of neuronal cells and reactivate mechanisms of neural differentiation and/or plasticity to achieve progression(57). Tumors may also be able to stimulate their innervation during cancer progression (58) and to invade already existing nerves along the perineural space(59). Both cancer cells and nerve fibers secrete factors that favor rapid growth of both, making the neural-epithelial interaction a mutually beneficial process(60). In addition, cancer cells themselves may acquire brain-like properties as an adaptation for brain colonization. For examples, i) breast-to-brain metastatic tissue and cells display phenotypes and metabolism similar to that of neuronal cells(61) and ii) malignant melanoma exhibits cytological characteristics of nerve cells(62). Interestingly, all the neuronal genes and the neuronal pathway coming up as strongly associated with *MAPT* expression were not detected in the P53 status analysis, suggesting that the association of *MAPT* with this neuronal pathway in several cancer types is independent of P53 status.

Genes modulated after *MAPT* KO in neuroblastoma cells (PAGANETTI_TAU_KO_VS_WT) were associated with *MAPT* expression in multiple cancer types. Importantly, genes downregulated after *MAPT* KO had a positive association with *MAPT* in brain tumors (GBM, PCPG, LGG) and vice versa for upregulated genes. Thus, gene modulation after *MAPT* KO in the neuroblastoma cell line was recapitulated in the *MAPT*-gene correlation analysis in brain cancer clinical samples. This supports the relevance of the pre-clinical model and the potential of our analysis to identify biological networks linked to *MAPT* expression.

Considering our recent findings(16), we aimed at characterizing the possible interplay between *MAPT* and P53 in cancer. *MAPT* expression levels were different according to P53 status in some cancer types. In BRCA, the lower *MAPT* expression observed in P53 mutant tumors may be explained by the fact that P53 mutations are found more in ER-, PR-basal carcinoma (∼88%) when compared to ER+, PR+ luminal tumors (∼26%) in which *MAPT* is upregulated by ER/PR(40). A negative association between *MAPT* expression and expression of P53 targets genes was detected in many cancer types, particularly in brain tumors. This agreed with the data obtained in neuroblastoma cells depleted of Tau (**Figure S1b**). However, a positive association was found in BRCA, KIRP, and UVM, unveiling some tissue specificity in this relationship.

Furthermore, we stratified the pan-cancer *MAPT*-gene correlation analyses by P53 status (i.e. mutated o WT) and revealed that the association of *MAPT* expression with cell cycle, inflammation, and EMT varies not only according to the cancer type but, in some instances according to P53 status. Additional hints on the connection between Tau and P53 came from the *MAPT*-*MDM2* correlation pattern. *MDM2* is the main E3 protein ubiquitin ligase and antagonist of P53 and had a high delta correlation in READ, driven by a negative correlation with *MAPT* in P53 tumors.

Our study expanded a previous report(19), based on an older version of the TCGA dataset, on the association of *MAPT* expression with patient survival. In that study, a positive correlation between *MAPT* expression and survival in glioma, breast cancer, kidney clear cell carcinoma, lung adenocarcinoma, and pheochromocytoma/paraganglioma was described. While the log-rank test was applied in the previous study(19), our analytical approach was based on the Cox regression model that we deemed more appropriate when running univariate and multivariate breakdowns. Doing so, we also identified a negative correlation in uterine cancer in a multivariate analysis. Moreover, we also separated low-grade glioma from glioblastoma to reveal a positive correlation only in the former, thereby providing additional information compared to previous studies grouping these two cancer types(63). We found a positive correlation between *MAPT* expression and survival in breast cancer, in line with previous studies(64–68). The association remained significant in the multivariate analysis, indicating independence from other prognostic factors. Estrogen and progesterone directly modulate the *MAPT* promoter(38, 69), explaining our observation of a strong positive correlation of *MAPT* with estrogen pathways and partially explaining the association with better survival.

The observation that *MAPT* is associated positively or negatively with survival depending on the cancer type, led us to the hypothesis that this could be linked with the distinct biological networks we found *MAPT* associated to. While a complex picture emerged, we identified genes for which their correlation with *MAPT* and *MAPT* hazard ratio were associated, partially confirming our hypothesis. Most of these genes encode either for structural proteins (i.e. COL5A3), components of the extracellular matrix (THSD4), or contributing to protein folding and/or degradation (FBXL16, PDIA6, PPIB). They are involved in proliferation, EMT, adhesion, and migration processes(43–47), thereby strengthening the observed association between *MAPT* expression and proliferation and EMT processes. Interestingly, the protein encoded by the best-associated gene, FBXL16, stabilizes C-MYC by blocking its ubiquitination and subsequently promotes cancer cell proliferation and migration(70).

In our characterization of *MAPT’s* role in cancer, we also investigated its association with drug response in cell lines derived from 22 different cancer types. The best-described association between Tau and drug response was with drugs targeting microtubules, such as taxanes. As a microtubule-binding protein, it has been proposed that Tau could compete with taxanes in the binding with microtubules. Low Tau is associated with a better response to taxanes in breast(69) ovarian(71, 72), gastric(73), prostate(74), and non-small-cell lung cancer(75). Nevertheless, some studies came to the opposite conclusion and some Paclitaxel trials did not confirm the predictive value of Tau determination(76–78). Our study did not detect a strong association between *MAPT* expression and microtubule-targeting drugs. Negative correlations were detected with Paclitaxel in gastric, plasma cell, and lung-derived lines, with podophyllotoxin in bone and pancreas-derived cells whereas a positive correlation is found with filanesib in liver-derived cells. Our study highlights, however, a strong association of *MAPT* expression with two other classes of cancer drugs, HDAC inhibitors, and several kinase inhibitors. A recent study suggests a possible predictive role of Tau expression in regards to HDAC inhibitors(79), however, the association of *MAPT* with sensitivity to kinases (PI3K, Aurora) inhibitors in bone-derived cells is a new finding. Osteosarcoma is a frequent type of pediatric bone cancer for which, despite many advancements in diagnostic technology, no efficient therapy approach has been identified because of the high metastasis rate and drug resistance(80). PI3K and Aurora kinase, both involved in mitosis and cell proliferation, are overexpressed in osteosarcoma and represent promising targets for osteosarcoma treatment(81, 82). The identification of *MAPT* as a predictive marker for response to these inhibitors may open, once validated in human biopsies, a new therapeutic perspective. We also detected strong positive and negative correlations between *MAPT* expression and drug responses in uterus-derived cell lines. Positive correlations are mainly detected with chemotherapy agents, whereas negative correlations are detected for Aurora and PI3K inhibitors, for example. Aurora kinase is frequently overexpressed in ovarian cancer and its expression has a prognostic value. Therefore the Aurora kinase family has evolved as a potential target for precision medicine in cancer(83). In the last decade, several Aurora kinase inhibitors have been developed and tested in several cancer types. Two recent clinical trials with different Aurora kinase inhibitors have shown activity in epithelial ovarian and clear-cell ovarian cancer(83). We may speculate that Tau interferes with Aurora kinase both at the level of the microtubule network and also at the level of P53, as multiple pieces of evidence underly a crosstalk between Aurora kinase and P53(83). The presence of several correlations between high *MAPT* expression and response to drugs with various modes of action in bone and uterus-derived cells indicate that *MAPT* expression may represent a powerful marker to predict response to combination therapies in bone and uterine cancer. Strikingly, a positive correlation between *MAPT* and response to several MEK inhibitors is found in breast and lung-derived lines. This could be particularly relevant in lung cancer where MEK inhibitors in combination with chemotherapy are highly significant for improving clinical efficacy and causing a delay in the occurrence of drug resistance(84).

A possible limitation of our study is that correlation-based analyses cannot imply causation. Experiments would be required to unveil the detailed molecular mechanisms, but this would be possible only in one or a few pre-clinical models, with known limitations in being representative of clinical tumors. On the contrary, in our analysis we were able to interrogate the largest molecularly characterized pan-cancer cohorts that, with over 10000 clinical specimens from 32 distinct cancer types and over 1000 pre-clinical samples, could better recapitulate the clinical disease and clinical heterogeneity, offering the opportunity to gain a comprehensive overview of the putative role of Tau in cancer, fostering and providing guidance for further research and validations.

Altogether, we described how several pathways and genes are associated with *MAPT* expression in all main cancer types. We comprehensively assessed the association between *MAPT* and survival, and the possible link with P53 status, and we present evidence that *MAPT* expression may affect the response to multiple classes of therapeutics.

## Materials and methods

### TCGA Dataset analysis

The Cancer Genome Atlas pan-cancer dataset was downloaded at https://gdc.cancer.gov/about-data/publications/pancanatlas (Downloaded files: EBPlusPlusAdjustPANCAN_IlluminaHiSeq_RNASeqV2.geneExp.tsv and clinical_PANCAN_patient_with_followup.tsv. A total of 11069 samples from 32 different cancer types were present in the dataset. The ‘acronym’ indicating the cancer type was not available for 213 samples that were excluded from the present study. Only genes with log10(FPKM+1) average expression above 0.5 and standard deviation above 0.2 in at least one cancer type were kept for downstream analysis (n= 17646). Tau expression was correlated with the expression of all genes using Spearman correlation, as implemented in the stats (v. 3.5.0) R base package.

P53 mutations and CNA status also downloaded from the GDC portal were grouped to identify WT and MUT samples and to distinguish functionally distinct groups according to the scheme in Table S2.

**Supplementary Table 2.**
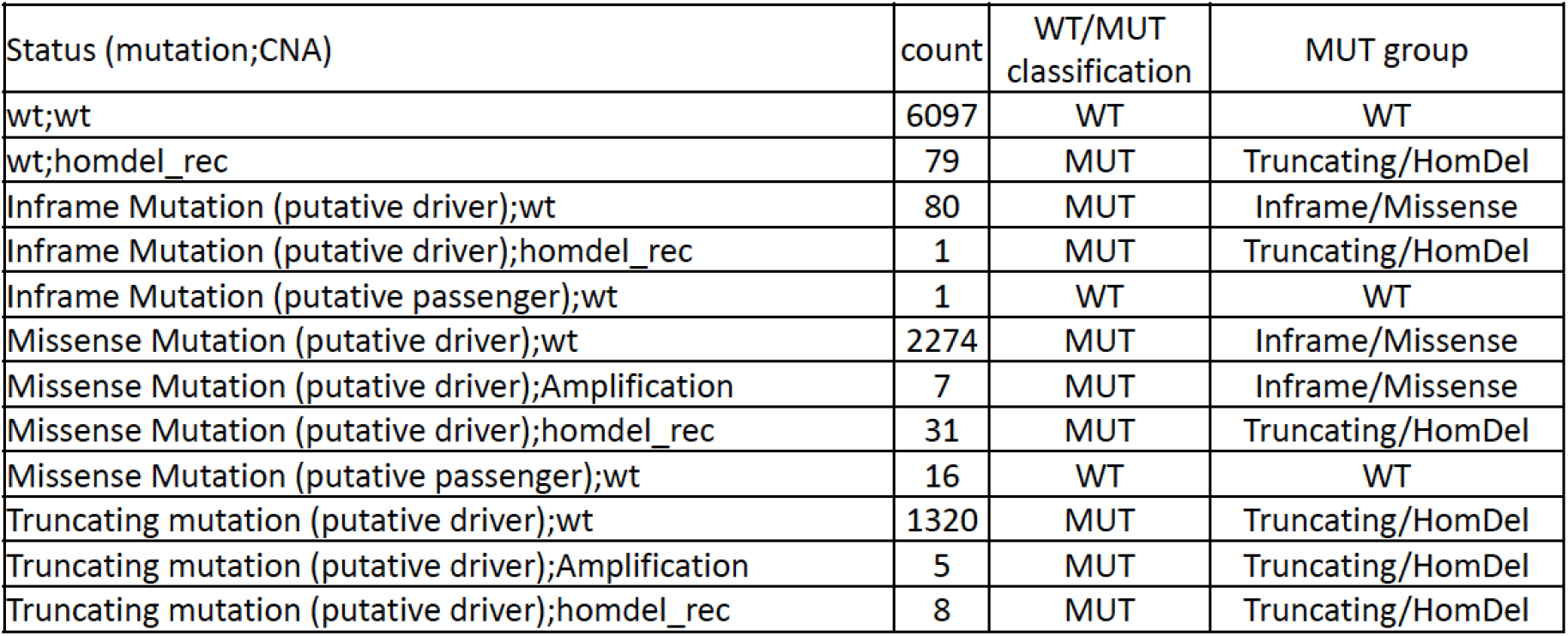

| Status (mutation;CNA) | count | WT/MUT classification | MUT group |
| --- | --- | --- | --- |
| wt;wt | 6097 | WT | WT |
| wt;homdel_rec | 79 | MUT | Truncating/HomDel |
| Inframe Mutation (putative driver);wt | 80 | MUT | Inframe/Missense |
| Inframe Mutation (putative driver);homdel_rec | 1 | MUT | Truncating/HomDel |
| Inframe Mutation (putative passenger);wt | 1 | WT | WT |
| Missense Mutation (putative driver);wt | 2274 | MUT | Inframe/Missense |
| Missense Mutation (putative driver);Amplification | 7 | MUT | Inframe/Missense |
| Missense Mutation (putative driver);homdel_rec | 31 | MUT | Truncating/HomDel |
| Missense Mutation (putative passenger);wt | 16 | WT | WT |
| Truncating mutation (putative driver);wt | 1320 | MUT | Truncating/HomDel |
| Truncating mutation (putative driver);Amplification | 5 | MUT | Truncating/HomDel |
| Truncating mutation (putative driver);homdel_rec | 8 | MUT | Truncating/HomDel |

### DEPMAP Dataset analysis

*MAPT* CRISPR KO, gene expression, proteomic and drug-response pan-cancer cell line data from the Dependency Map initiative were downloaded from https://depmap.org/portal/. (Downloaded files from version 21Q1: CCLE_expression.tsv, protein_quant_current_normalized.csv, sanger-dose-response.csv, sample_info.csv; from PRISM Repurposing 19Q4: primary-screen-replicate-collapsed-logfold-change.csv) Cancer types with more than one cell line present in the gene expression dataset were included. Cell lines labelled as “Engineered” and “Fibroblast” were excluded. After this filtering, gene expression was available for 1320 cell lines from 28 cancer types. Drug response data were available for 692 of them, belonging to 22 cancer types. The area under the drug response curve (AUC), ranging between 0 (response) and 1 (resistance) was correlated with Tau expression for each cancer type. Spearman correlation was performed only when at least 10 cell lines were tested and at least two sensitives (AUC<0.8) and two resistant (AUC>0.8) were present. A total of 121 compounds tested in at least 11 of 22 cancer types were reported. Normalized mass spectrometry data across 375 cell lines(85) were used to evaluate the correlation between *MAPT* gene expression and Tau protein levels. All datasets after sample and/or feature filtering are made available as described in the Data availability section.

### *MAPT* knock-out signature in neuroblastoma cells

RNA-Seq data derived from human neuroblastoma SH-SY5Y wild-type or KO for Tau (accession no. E-MTAB-8166) were downloaded from (https://www.ebi.ac.uk/biostudies/arrayexpress/studies/E-MTAB-8166?key=64a67428-adb9-4681-99c9-98910b78ed4c). The top 100 genes upregulated and the top 100 downregulated after *MAPT* KO generated two genesets (PAGANETTI_TAU_KO_VS_WT_UP and PAGANETTI_TAU_KO_VS_WT_DOWN respectively) included in the GSEA.

### Statistical analyses

All statistical analyses were performed using R (v 4.2, platform: x86_64-apple-darwin17.0, running under macOS Big Sur 11.4). Association between two continuous variables (i.e. gene-gene expression) was quantified by Spearman’s correlation analysis. Association with overall survival was performed as univariate or multivariate Cox regression analysis as implemented in the *survival* (v.3.1) R/Bioconductor package. The analysis was performed when at least 10 events (deaths) were present in the considered subset of cancer cases and when, in multivariate analysis, information for at least three of the four covariates (Size, Lymph node status, Metastatic status, AURKA gene expression) was available in at least 20 patients. P-value < 0.05 indicate a significant association. *MAPT* expression was dichotomized in high or low using the median expression as a cut-off.

### GeneSet Enrichment Analysis (GSEA)

A custom list of genesets to be tested was created as follows. The HALLMARK geneset collection was downloaded from the MSigDB (https://www.gsea-msigdb.org/gsea/msigdb/, v 7.1). The genesets related to the PANTHER classification system(86) were downloaded from https://maayanlab.cloud/Harmonizome/dataset/PANTHER+Pathways. Four further genesets were added, three related to senescence (SENESCENCE_HERNANDEZ-SEGURA, SENESCENCE_PURCELL, SENESCENCE_CASELLA)(26, 27, 29), one collecting P53 direct targets (PAGANETTI_TP53_DIRECT_TARGET) (30–33) and PAGANETTI_TAU_WT_VS_KO described above. Correlation-ranked genes were tested for geneset enrichment using the gsea function from the *phenoTest* (v.1.28.0) R/Bioconductor package. In the analysis stratified by P53 status, an absolute delta NES between P53 mutant and WT tumors was considered significant.

Selected PANTHER pathways were represented and overlaid with MAPT-gene correlation values in selected cancer types using the SBGNview (v. 1.10.0) R/Bioconductor package(87).

### Identification of top 100 genes co-express with MAPT using EnrichR

Using EnrichR(88) (https://maayanlab.cloud/Enrichr/) we identified the top 100 genes co-expressed with MAPT in the ARCHS4 human tissue RNA-seq dataset. Genes were overlapped with our list of 809 genes that we found correlated with MAPT in at least one cancer type.

## Acknowledgment

We thank all members of the Aging Disorders Laboratory for their support and advice during this study. This work was generously supported by the Gelu Foundation, the Mecri Foundation, and the Gabriele Foundation. We thank PD Dr. Andrea Rinaldi, Genomics Facility BIOS+, Bellinzona Switzerland, for generating the RNAseq data.

## Authors contributions

Maurizio Callari performed the bioinformatic analysis, collected the data, prepared figures, and contributed to the manuscript writing; Martina Sola and Claudia Magrin contributed to the interpretation of the data; Andrea Rinaldi produced the cDNA library and performed RNAseq analysis; Marco Bolis processed and analyzed the RNAseq raw data; Luca Colnaghi contributed to the concept and overview of the study; Paolo Paganetti contributed to the concept and overview of the study and provided the financial support, Stéphanie Papin conceived the study, interpreted the data, and wrote the first manuscript draft. All authors revised the final manuscript version.

## Code availability

No custom algorithms have been developed in this study. Reference code related to the main analyses performed is available at https://github.com/mauricallari/MAPT

## Data availability

Imported and filtered datasets used to generate the presented results are available on Zenodo (DOI: 10.5281/zenodo.8069665, https://zenodo.org/record/8069665).

