## Supplementary figures and images for "Cancer-specific association between Tau (*MAPT*) and cellular pathways, clinical outcome, and drug response"

### Supplementary Figure 2

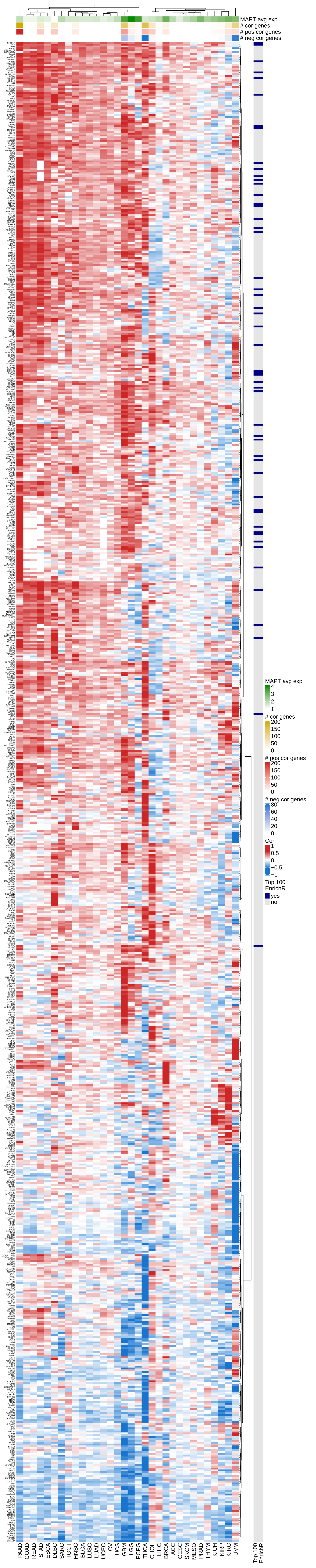

### Supplementary Figure 6

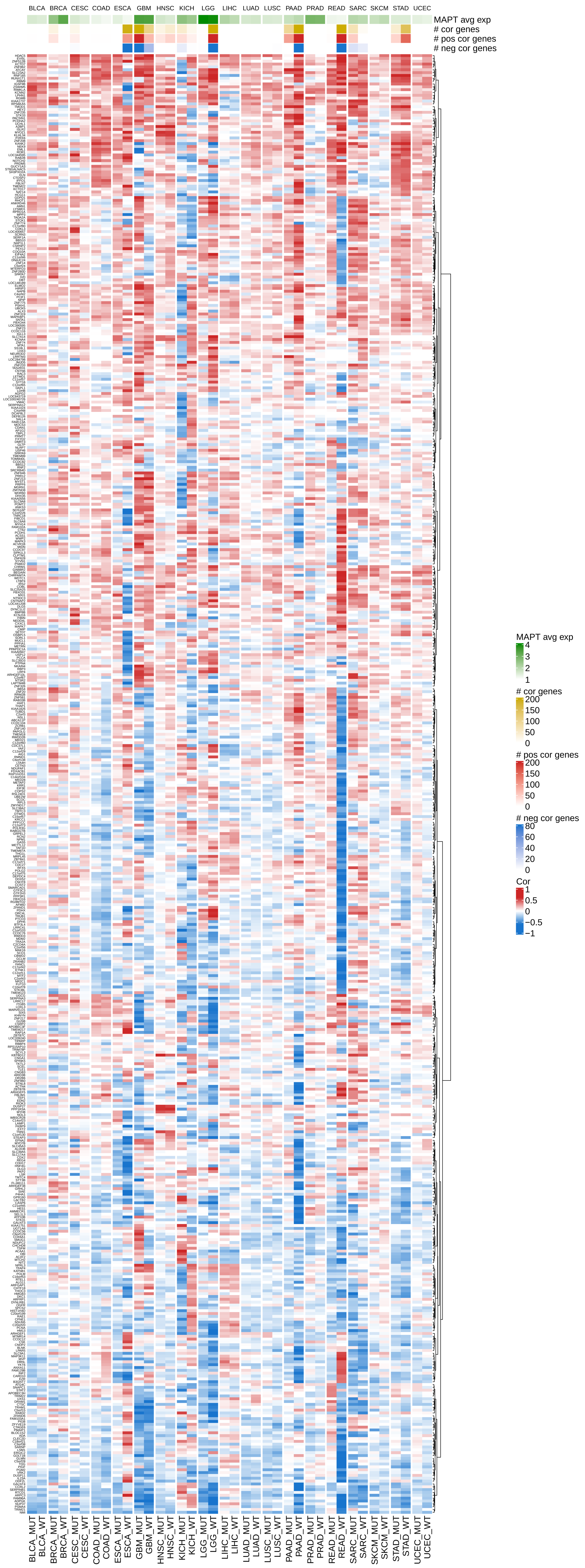

### Supplementary Figure 7

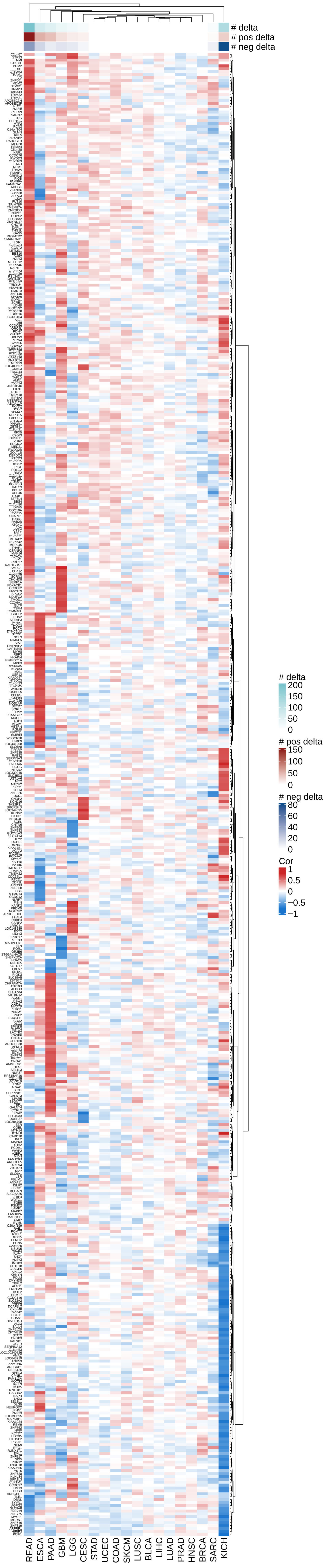

### Supplementary Figure 8

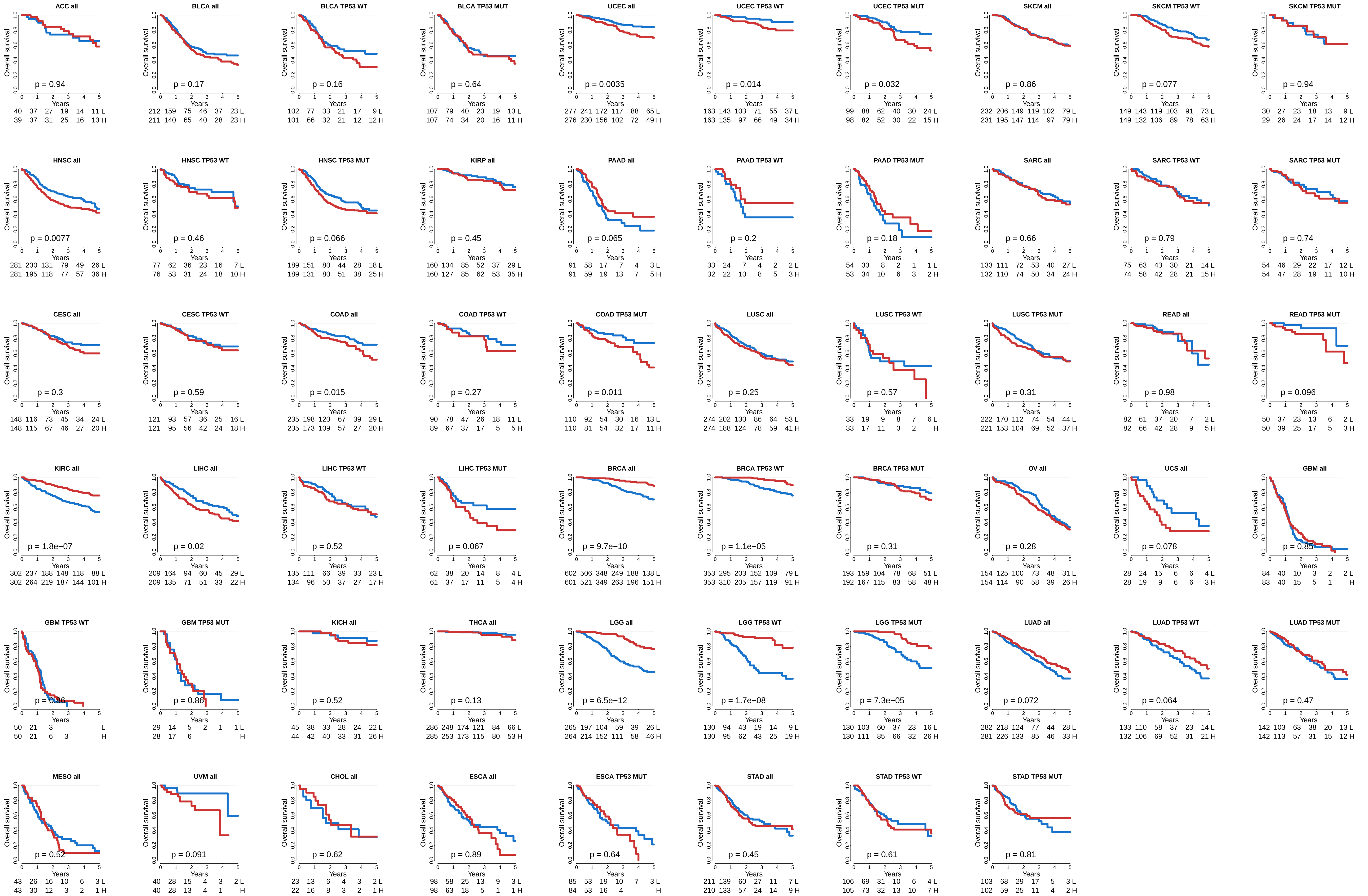
